# Decoding herbal materials of representative TCM preparations with the multi-barcoding approach

**DOI:** 10.1101/2020.06.29.177188

**Authors:** Qi Yao, Xue Zhu, Maozhen Han, Chaoyun Chen, Wei Li, Hong Bai, Kang Ning

**Author notes:** These authors contributed equally to this work (Qi Yao, Xue Zhu).

## Abstract

With the rapid development of high-throughput sequencing (HTS) technology, the techniques for the assessment of biological ingredients in Traditional Chinese Medicine (TCM) preparations have also advanced. By using HTS together with the multi-barcoding approach, all biological ingredients could be identified from TCM preparations in theory, as long as their DNA is present. The biological ingredients of a handful of classical TCM preparations were analyzed successfully based on this approach in previous studies. However, the universality, sensitivity and reliability of this approach used on TCM preparations remain unclear. Here, four representative TCM preparations, namely Bazhen Yimu Wan, Da Huoluo Wan, Niuhuang Jiangya Wan and You Gui Wan, were selected for concrete assessment of this approach. We have successfully detected from 77.8% to 100% prescribed herbal materials based on both ITS2 and *trnL* biomarkers. The results based on ITS2 have also shown a higher level of reliability than those of *trnL* at species level, and the integration of both biomarkers could provide higher sensitivity and reliability. In the omics big-data era, this study has undoubtedly made one step forward for the multi-barcoding approach for prescribed herbal materials analysis of TCM preparation, towards better digitization and modernization of drug quality control.

## 1. Introduction

Traditional Chinese Medicine (TCM) preparation has been used in clinics in China for at least 3,000 years^1,2^. It has been utilized to prevent and cure various diseases in China and has become more popular all over the world during the last decades. TCM preparation is composed of numerous plants, animal-derived and mineral materials. According to the guidance of Chinese medicine theory and Chinese Pharmacopeia (ChP)^3^, different medicinal materials were crushed into powder or boiled, then mixed and molded into pills together with honey or water to get a TCM preparation (also called patented drug). Although TCM preparations have been extensively used in recent years, many problems remain to be resolved, such as quality control (QC), in which particular attention should be focused on its materials and production process to ensure its efficacy and safety. The quality of TCM preparations is the prerequisite for their clinical efficacy, its quality assessment includes the qualitative and quantitative analysis of chemical ingredients and biological ingredients^4^. Current methods for the QC of TCMs have been mainly assessed based on chemical profiling^4^ (e.g. TLC^5^, HPLC-UV^6,7^, HPLC-MS^8^). Through comparing with reference herbal materials or targeted compounds, TLC and HPLC method can retrieve species information but not precise enough, especially in identifying the hybrid species of genetics, which might occur the incorrect identification, introduce biological pollution and adulteration during the herbal materials collection and preprocessing. However, the utilization of DNA, a fragment that stably exists in all tissues^9^, could identify herbal materials at species level accurately, providing a higher level of sensitivity and reliability, thus complementing the drawback of chemical analysis^10,11^.

The concept of biological ingredient analysis based on DNA-barcoding was proposed by Hebert^12^. Chen *et al*. have first applied a serval candidate DNA barcodes to identify medicinal plants and their closely related species^13^. Coghlan *et al*., for the first time, have used DNA barcoding to determine whether TCM preparations contain derivatives of endangered, trade-restricted species of plants and animals^2^. In 2014, Cheng *et al*. have first reported the biological ingredients analysis for Liuwei Dihuang Wan (LDW) using the metagenomic-based method (M-TCM) based on ITS2 and *trnL* biomarkers^14^. After that, the reports on the herbs of TCM preparations based on DNA biomarkers have been sprung up, such as Yimu Wan^15^ (YMW), Longdan Xiegan Wan^16^ (LXW) and Jiuwei Qianghuo Wan^17^ (JQW). Interestingly, recent studies have reported several TCM preparations that might be effective in the prevention and treatment for COVID-19^18,19^, such as Lianhua Qingwen capsule^20^, Jinhua Qinggan granules^20^, Yiqi Qingjie herbal compound^21^, etc. Among these, Lianhua Qingwen capsule, is reported to be effective in the prevention or treatment for COVID-19 mainly due to its biological ingredients such as Glycyrrhizae Radix Et Rhizoma and Rhei Radix Et Rhizome^3^. The same principle applies for Jinhua Qinggan granules and Yiqi Qingjie herbal compound. These findings again emphasized the importance of biological ingredient analysis of TCM preparations.

A TCM preparation can be regarded as a “synthesized mixture of species”, which resembles the analytical target of metagenomic approach. In metagenomics approach, based on suitable DNA biomarkers, the genetic information of all DNA-contained ingredients could be obtained in a most effective and cost-effective way via HTS. Due to the conservation of ITS2^22^ and its high inter-specific and intra-specific divergence power^23–25^, and the convenience of amplification DNAs from heavily degraded samples based on a short fragment *trnL*^26–28^, these two fragments are usually chosen as biomarkers for herbal species identification. Such an approach based on multiple barcodes for herbal ingredient analysis is referred to as the “multi-barcoding approach”.

In spite of scientific advances of recent studies, the solidity (*i.e*., universality, sensitivity and reliability) of multi-barcoding approach on identifying a variety of biological ingredients of TCM preparations simultaneously remains unclear and needs to be investigated systematically. Therefore, we selected three TCM preparations with simple compositions and pervasively used named Niuhuang Jiangya Wan (NJW), Bazhen Yimu Wan (BYW), Yougui Wan (YGW), and one TCM preparation named Da Huoluo Wan (DHW) with much more complicated components and widely applied, as targets for herbal materials assessment by using ITS2 and *trnL* biomarkers. Based on the assessment of their prescribed herbal species (PHS) of the prescribed herbal materials (PHMs), the universality, sensitivity and reliability of the multi-barcoding approach have been evaluated, from which the multi-barcoding approach stands out as a superior method for PHMs analysis for TCM preparations.

## 2. Materials and Methods

### 2.1. Sample collections

Four TCM preparations, each purchased from two different manufacturers (marked as A and B) with three batches (I, II and III), were collected (**Supplementary Table 1**). Each batch was implemented with three biological replicates based on ITS2 and *trnL* respectively. Therefore, 4*2*3*3*2=144 samples in total were used for the subsequent experiment. Here, we gave an example to clarify the mean of SampleID: DHW.A.I1 means the DHW sample was bought from the first batch of manufacturer A, and it was one of the three biological replicates (I1) of the first batch (I).

### 2.2. DNA extraction and quantification

For DNA extraction, we used an optimized cetyl trimethyl ammonium bromide (CTAB) method (TCM-CTAB)^29^. Each sample (1.0 g) was completely dissolved with 0.1 M Tris-HCl, 20 mM EDTA (pH 8.0, 2 ml). Dissolved solution (0.4 mL) was diluted with extraction buffer (0.8 mL) consisting of 2% CTAB; 0.1 M Tris-HC1 (pH 8.0); 20 mM EDTA (pH 8.0); 1.4 M NaCl, and then 100 *μ*L 10% SDS, 10 *μ*L 10 mg/mL Proteinase K (Sigma, MO, USA) and 100 *μ*L *β*-Mercaptoethanol (Amresco, OH, USA) were added and incubated at 65 °C for 1 h with occasional swirling. Protein was removed by extracting twice with an equal volume of phenol: chloroform: isoamyl-alcohol (25: 24: 1), and once with chloroform: isoamyl-alcohol (24: 1). The supernatant was incubated at −20 °C with 0.6 folds of cold isopropanol for 30 min to precipitate DNA. The precipitate was washed with 75% ethanol, dissolved and diluted to 10 ng/*μ*L with TE buffer, and then used as a template for PCR amplification (**Supplementary Figure 1**). DNA concentration was quantified on Qubit^®^2.0 Fluorometer.

### 2.3. DNA amplification and DNA sequencing

The PCR amplification was performed in a 50 *μ*L reaction mixture that contain 1 *μ*L of DNA extracted from TCM preparations, 10.0 *μ*L of 5×PrimeSTAR buffer (Mg^2+^ plus) (TaKaRa), 2.5 *μ*L of 10 *μ*M dNTPs (TaKaRa), 0.5 *μ*L each of forward and reverse primers (10 μM), 2.5 *μ*L dimethylsulfoxide (DMSO) and 0.5 *μ*L PrimeSTAR^®^ HS DNA Polymerase (Takara, 2.5 U/*μ*L). For amplification and sequencing of ITS2 region, the forward primers S2F^13^ and the reverse primer ITS4^30^ (**Supplementary Table 2**) with seven bp MID tags (**Supplementary Table 3**) were designed for PCR amplification. PCR reactions were implemented as follows: pre-denaturation at 95 °C for five min, then 10 cycles made up of 95 °C for 30 s and 62 °C for 30 s with ramping of −1 °C per cycle, followed by 72 °C for 30 s, next followed by 40 cycles of 95 °C for 30 s, 55 °C for 30 s and 72 °C for 30 s; the procedure ended with 72 °C for 10 min. For *trnL* region, the forward primers *trnL-c* and the reverse primer *trnL-h* with 7 bp MID tags were also designed for PCR amplification. The PCR reactions were carried out according to the conditions: pre-denaturation at 95 °C for five min, 10 cycles made up of 95 °C for 30 s and 62 °C for 30 s with ramping of −1 °C per cycle, followed by 72 °C for 30 s; then followed by 40 cycles of 95 °C for 30 s, 58 °C for 30 s and 72 °C for 30 s; the procedure ended with 72 °C for 10 min. For better amplification effect, touchdown PCR^30,31^ was carried out. The PCR products were electrophoresed on 1% agarose gel (**Supplementary Figure 2**) and purified with QIAquick Gel Extraction kit (QIAGEN). The DNA concentration was quantified on Qubit 2.0 Fluorometer. After removing one *trnL*-marked BYW specimen that failed to be amplified, which was potentially caused by severe PCR inhibition, and one ITS2-marked YGW sample that failed to be built the next-generation sequencing library preparation, 142 samples (**Supplementary Table 4**) were sent for Illumina MiSeq PE300 paired-end sequencing. The raw sequencing data for TCM preparation samples were deposited to the NCBI SRA database with accession number **PRJNA562480**.

### 2.4. Sequencing data analysis procedure and software configuration

We first used the FastQC software (version 0.11.7) with default parameters to evaluate the quality of the sequencing reads. Reads from the same sample were assembled together by using QIIME script ‘join_paired_end.py’. Then we used the ‘extract_barcodes.py’ to extract the double-end barcodes from all reads, and the ‘split_libraries_fastq.py’ was used to split the sample according to their barcodes (**Supplementary Table 3**) from the mixed sequencing data, and we also used its ‘-q 20 --max_bad_run_length 3 --min_per_read_length_fraction 0.75 --max_barcode_errors 0 --barcode_type 7’ parameters to preliminarily filter the low-quality sequences, then Cutadapt software (version 1.14) was used to remove the primers (**Supplementary Table 2**) and adapter from all samples.

These reads of all samples were QCed by MOTHUR^32^ (version 1.41.0). Per reads of ITS2 whose length is <150 bp or >510 bp and the reads of *trnL* whose length is <75 bp were removed. After that, we discarded the sequence whose average quality score was below 20 in each five bp-window rolling along with the whole reads. Then the sequences that contained ambiguous base call (N), homopolymers of more than eight bases or primers mismatched, uncorrectable barcodes, were also removed from ITS2 and *trnL* datasets.

To match the target species for each sequence, we used the BLASTN (E-value=1E-10) to search in ITS2 and *trnL* database based on GenBank^33^, respectively. Among all results, we first chose the prescribed herbal species with the highest score, else we selected the top-scored species. In addition, we also manually searched all prescribed herbal species of prescribed herbal materials in all samples. Then, we discarded the corresponding species of ITS2 and *trnL* sequences with relative abundance below 0.002 and 0.001, respectively. Rarefaction analysis was performed with R^34^ (version 3.5.2) using the “vegan” package (https://cran.r-project.org/web/packages/vegan/index.html) to evaluate the sequencing depth of TCM preparations samples.

To understand the difference of samples between manufacturers and batches, the distance between any two samples was calculated based on Euclidean distance. By using the sample as the node and the distance of any two samples as the edge, we built a network cluster for each TCM preparation and visualized in Cytoscape^35^ (version 3.7.1) based on ITS2 and *trnL*, respectively. Principal component analysis (PCA) analysis was also utilized in R package “ade4” (https://cran.r-project.org/web/packages/ade4/index.html) to detect the difference between two manufacturers based on clustering result. We also used the LDA Effect Size (LEfSe)^36^ to select legacy biomarker, and then performed feature selection using minimum Redundancy Maximum Relevance Feature Selection (mRMR)^37^ to select the most discriminative biomarkers. The receiver operating characteristic (ROC) curve^38^ analysis was applied to evaluate the classification effectiveness of the biomarker selected from different manufacturers.

### 2.5. Terminology and abbreviation definitions

The prescribed herbal materials were defined as the herbal materials of a TCM preparation contained and recorded in ChP, abbreviated to PHMs.

The prescribed herbal species (abbreviated to PHS) was the original species of PHMs, any one of them should be considered as the PHS.

The species that has the same genus with PHS was defined as substituted herbal materials (SHS), the species excluded the two above species was considered as the contaminated herbal species (CHS).

For easier understanding of the abbreviations of used in this study, we took one TCM preparation, YGW, as an example, as shown in **Table 1**, and the information for other TCM preparations was shown in **Supplementary Table 5**. We have also provided detailed information about the animal and mineral materials for the four TCM preparations in **Supplementary Table 6**.

**Table 1.**
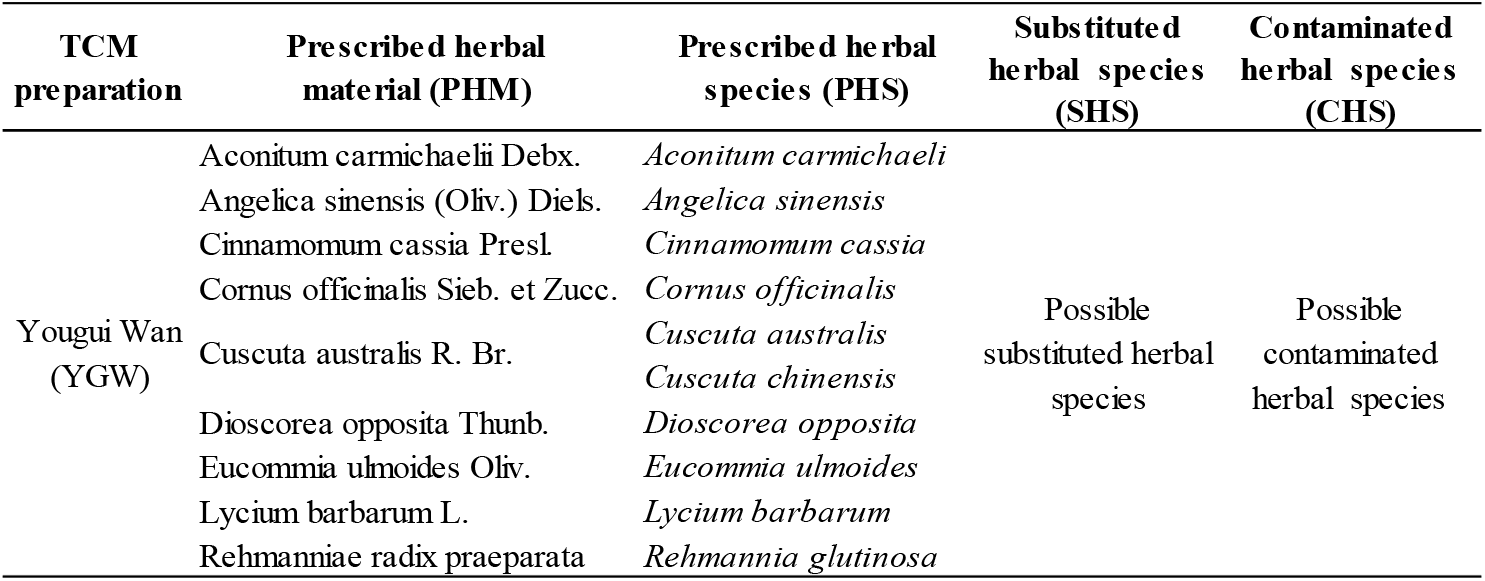
The abbreviation of prescribed herbal materials, un-prescribed herbal materials and its corresponding prescribed herbal species of Yougui Wan (YGW).

The universality was a measurement to evaluate how multi-barcoding approach could be applied on a broad scope of TCM preparations. The four representative TCM preparations were selected for this purpose.

The sensitivity was defined as the ratio of the number of detected PHMs, over the number of PHMs that could be identified in theory, that is,

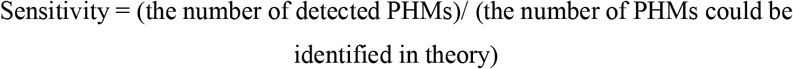

The reliability was defined as the number of detectable PHMs from the TCM preparations by multi-barcoding approach. The larger number of detectable PHMs, the better reliability.

## 3. Results

### 3.1. Profiling the prescribed herbal materials for all TCM preparations in Chinese pharmacopoeia

While several widely-used TCM preparations including LDW^14^, YMW^15^, LXW^16^ and JQW^17^, have their herbal materials assessed recently, it is important to choose representative preparations for deeper understanding of multi-barcoding approach for the quality assessment of TCM preparation. We examined the all ingredients (including herbs, animals and minerals) and herbal materials only for the TCM preparations recorded in ChP (2015 version) (**Supplementary Figure 3A** and **B**). On the basis of **Supplementary Figure 3A and B**, it is obvious that most TCM preparations have less than 25 ingredients (the total number of herbs, animals and minerals) and 20 herbal materials, respectively. Therefore, we selected three TCM preparations (BYW, NJW and YGW) with simple compositions and pervasively applications, and one TCM preparation (DHW) with much more complex ingredients, on the assessment of multi-barcoding approach for the quality assessment of TCM preparation. The detailed ingredients information about these four representative TCM preparations (such as the number of their herbal, animal and mineral materials) was shown in **Supplementary Figure 3C**.

### 3.2. Overview of the herbal materials from TCM preparations

The four selected TCM preparations were purchased from two manufacturers with three batches each. Each sample was implemented with three biological replicates based on ITS2 and *trnL*, respectively. Thus, 144 samples were obtained for DNA extraction, PCR amplification, library building. In this process, except for two failed samples, 142 samples were subjected to paired-end sequencing for subsequent data analysis.

After preliminary filtering (see more details in **Materials and Methods**), we obtained 25,271,042 ITS2 and 27,599,145 *trnL* sequencing reads, averages of 48,493 (BYW), 87,911 (DHW), 161,025 (NJW) and 58,501 (YGW) ITS2 sequencing reads per sample, respectively, and 57,954 (BYW), 139,512 (DHW), 129,560 (NJW) and 61,685 (YGW) *trnL* sequencing reads per sample, respectively (**Table 2**). Then rarefaction analysis was performed for each sample to detect whether the sequencing depth enough. At around 10,000 sequences per sample, all curves tended to approach the saturation plateau, suggesting that the sequencing depth was enough to capture all species information in all samples for the four TCM preparations (**Supplementary Figure 4**). Considering the smaller *trnL* database comparing to the database of ITS2, we filtered the corresponding species of ITS2 and *trnL* sequences with the relative abundance below 0.002 and 0.001, respectively. There were 47,533, 86,422, 160,712 and 58,008 remained on average of per sample of BYW, DHW, NJW and YGW based on ITS2, and 56,367 (BYW), 130,330 (DHW), 129,012 (NJW) and 59,709 (YGW) based on *trnL* (**Table 2**).

**Table 2.** The average number of reads of each sample after preliminary QC and threshold filtration for the four TCM preparations.

|  | **Biomarker** | **BYW** | **DHW** | **NJW** | **YGW** |
| --- | --- | --- | --- | --- | --- |
| Preliminary QC | ITS2(150~510 bp) | 48,493 | 87,911 | 161,025 | 58,501 |
| | *trnL*(≥ 75 bp) | 57,954 | 139,521 | 129,560 | 61,685 |
| Threshold selection | ITS2(≥ 0.002) | 47,553 | 86,642 | 160,712 | 58,008 |
| | *trnL*(≥ 0.001) | 56,367 | 130,330 | 129,012 | 59,709 |

Note that, QC means quality control, the reads were removed that below 150 bp or over 510 bp for ITS2, and the reads less than 75 bp for *trnL*, or the sequences that had an average quality score < 20 in each 5 bp-window rolling along with the whole read. Then we filtered out the sequences whose corresponding species was evidenced by the relative abundance less than 0.002 for ITS2 and 0.001 for *trnL*.

In general, several herbal materials have more than one prescribed herbal species, such as licorice, recorded as Glycyrrhizae Radix Et Rhizoma in ChP, includes species of *Glycyrrhiza uralensis, Glycyrrhiza inflate* and *Glycyrrhiza glabra*. Consequently, anyone original species of prescribed herbal materials (PHMs) should be regarded as prescribed herbal species (PHS). In this work, BYW contained eight prescribed herbal materials, NJW and YGW contain nine PHMs, and DHW contains 36 PHMs, they include 11, 15, 10 and 57 PHS (listed in **Supplementary Table 5** and **Table 1**), respectively.

The results of the ITS2 audit on 18 BYW samples, average of 8.2 PHS, 1.0 substituted herbal species (SHS, the species has the same genus with PHS), and 13.8 contaminated herbal species (CHS, the other detected species expect PHS and SHS) was detected, while 5.0 PHS, 0.3 SHS and 14.9 CHS were found in each *trnL* samples (**Figure 1A and B**). For DHW, each sample has the average of 23.7 PHS, 5.1SHS and 21.1 CHS based on ITS2, while average of 17.9 PHS, 6.8 SHS and 27.7 CHS based on *trnL* (**Figure 1C and D**). For NJW samples, average of 7.2 PHS, 2.8 SHS and 1.8 CHS were detected in individual samples based on ITS2, which was more than *trnL* (3.0 PHS, 3.0 SHS and 24.0 CHS; **Figure 1E and F**). The mean values of PHS, SHS and CHS detected in per YGW sample were 4.8, 0.9 and 10.4, and 3.7, 0.5, 17.3 based on ITS2 and *trnL*, respectively (**Figure 1G and H**). These differences may partially be due to the completeness of ITS2 and *trnL* database, as well as their intrinsic resolution properties.

**Figure 1.**
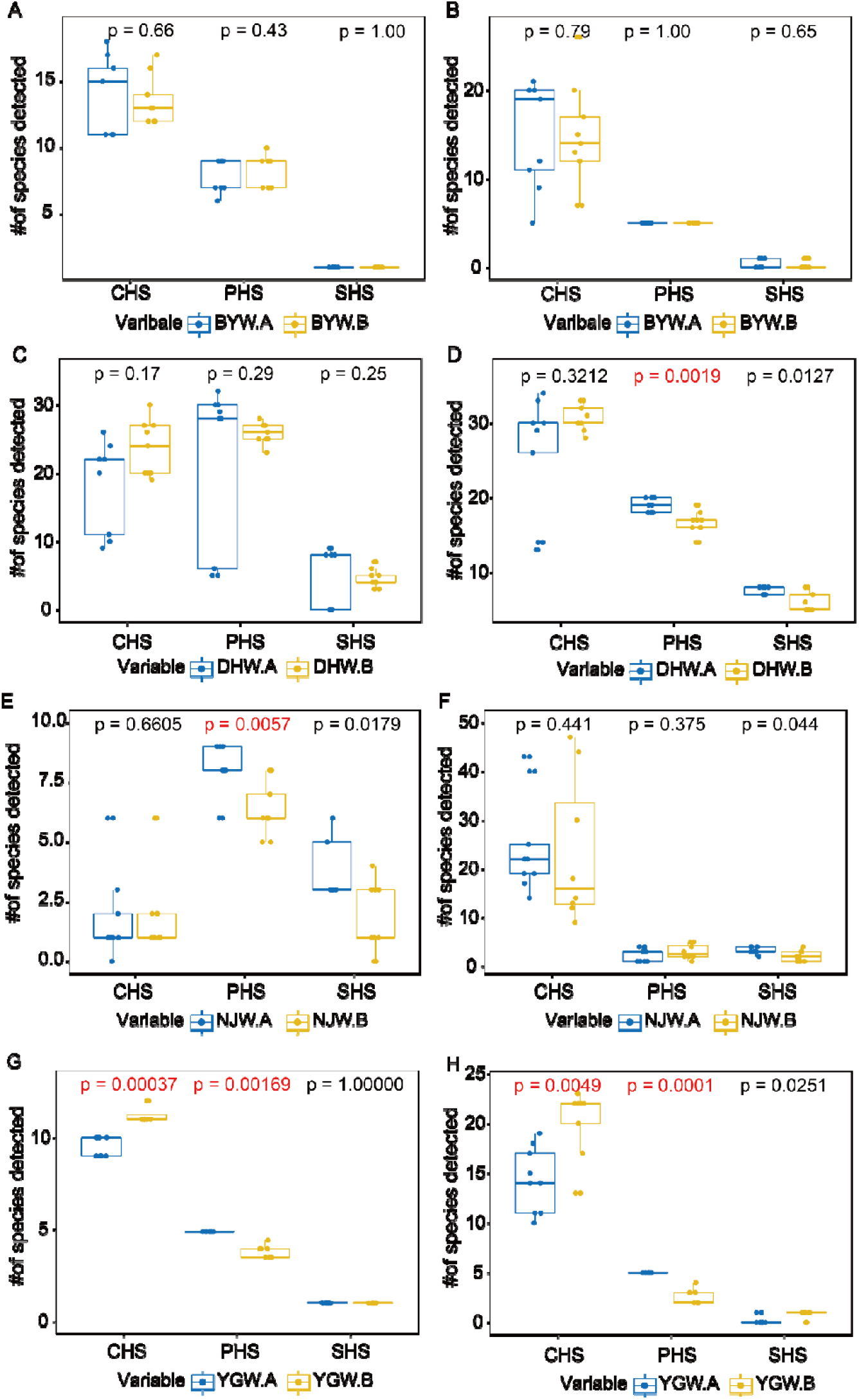
The number of detected prescribed herbal species (PHS), substituted herbal species (SHS) and contaminated herbal species (CHS), of all samples for per TCM preparation from two manufacturers (A & B). (A) BYW samples based on ITS2; (B) BYW samples based on *trnL*; (C) DHW samples based on ITS2; (D) DHW samples based on *trnL*; (E) NJW samples based on ITS2; (F) NJW samples based on *trnL*. (G) YGW samples based on ITS2; (H) YGW samples based on *trnL*.

The phylogenetic trees for each sample were also built based on the ITS2 and *trnL* datasets (**Figure 2** for DHW samples and **Supplementary Figure 5** for other TCM preparations). Each species whose relative abundance was greater than or equal to 0.1%, was displayed with 100% resolution in this tree (that is, any species existed in a sample could be exactly identified at species level). The genetic relationship and the coverage of the detected species were scattered widely, indicating the high sensitivity of the designed primer, and also confirmed there was no biological bias in our experiment.

**Figure 2.**
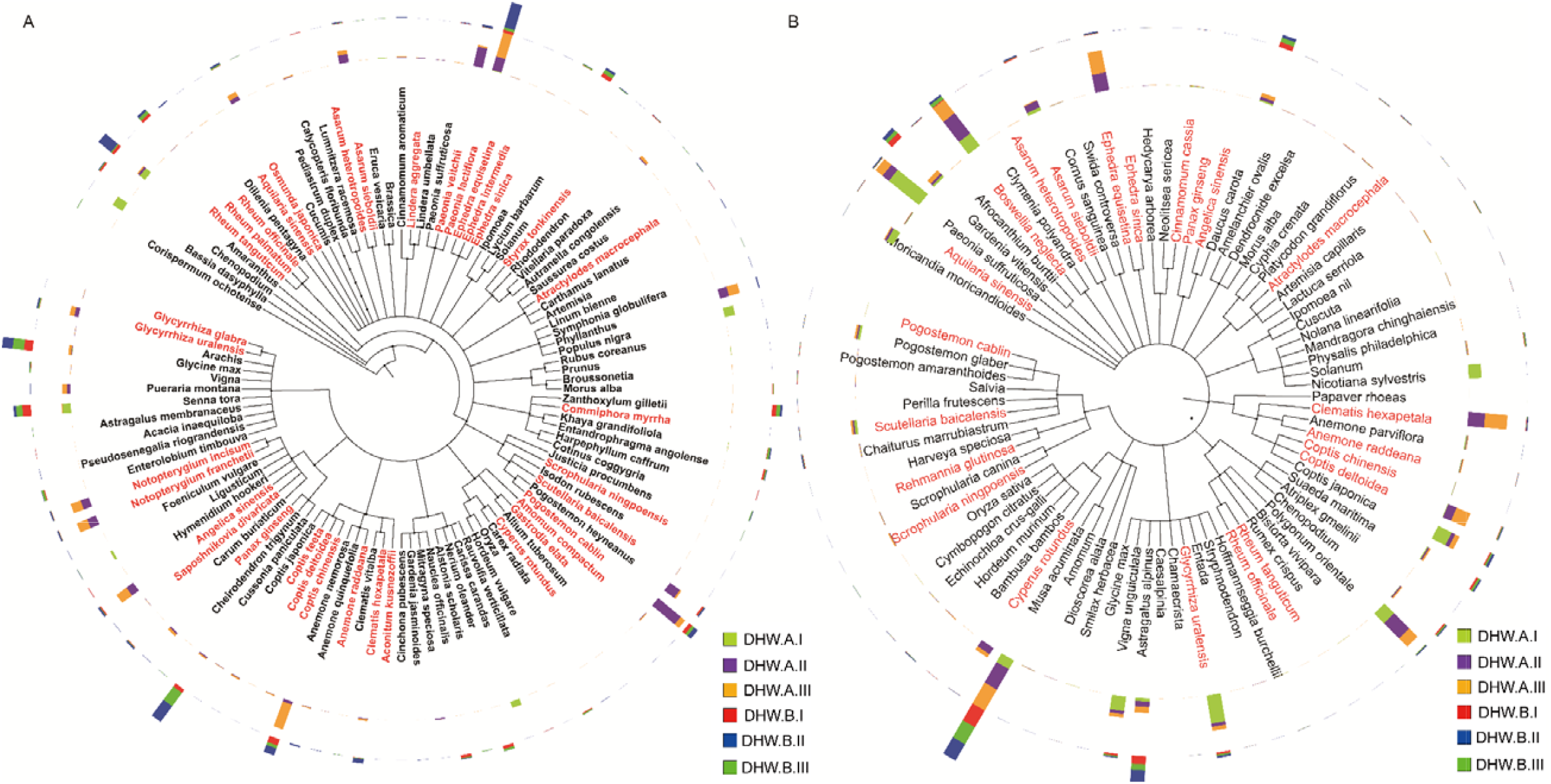
Phylogenetic analysis of the representative species that had at least 0.1% relative abundance for DHW samples. (A) Based on ITS2; (B) Based on *trnL*. The branch depicts the taxonomic classification of species. The word marked in red means the prescribed herbal species, and the colorful bar mea**ns** the average relative abundance of species across the three batches from the two manufacturers (A&B).

In summary, the multi-barcoding approach could accurately identify the herbal materials, including prescribed, substituted, and contaminated materials, for representative TCM preparations (including BYW, DHW, NJW and YGW). The result has demonstrated that the multi-barcoding approach has good universality for detecting PHMs from TCM preparation samples.

### 3.3. Sensitivity analysis of on prescribed herbal materials from TCM preparations

For more detailed probing of the composition of TCM preparations, we chose one TCM preparation named NJW with a relatively simple composition and pervasively application, and another TCM (DHW) with more complex ingredients, as targets to decode their PHMs through identifying their prescribed herbal species of each TCM preparations based on ITS2 and *trnL* datasets.

#### Analysis of herbal materials in the TCM preparations based on ITS2

The result of the ITS2 auditing on NJW samples, revealed that it could successfully detect all PHMs (9 herbal materials), including the processed herbal materials (such as the extractive of Huangqin), covering 12 detected PHS (**Table 3**). *Senna obtusifolia* (the average relative abundance was 48.4%) and *Senna tora* (45.4%) were the dominant species in all samples, followed by *Paeonia lactiflora* (3.4%) and *Ligusticum chuanxiong* (1.0%), suggesting that the modified CTAB method was suitable to extract their DNA and the primers were more suitable to amply their sequences. Besides the prescribed herbal species, seven substituted herbal species were also found, belonging to *Codonopsis, Ligusticum, Mentha, Paeonia* and *Senna* (their average relative abundance was 0.035%) and six possible contaminated genera namely *Ipomoea, Amaranthus, Anemone, Cuscuta, Pogostemon* and *Zanthoxylum*, which might be introduced during the biological experiment.

**Table 3.**
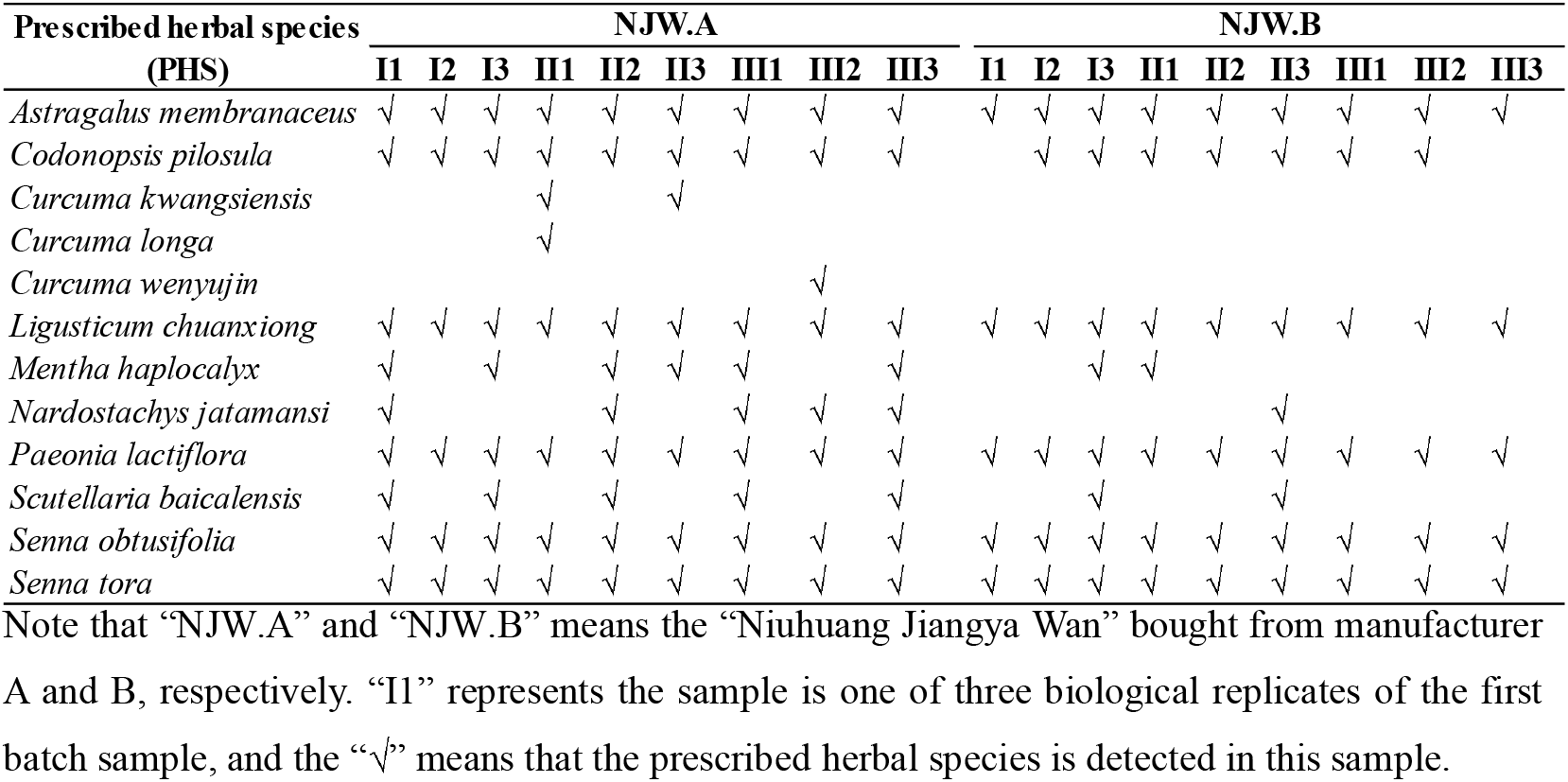
Prescribed herbal species for NJW preparation and their presence in each sample by multi-barcoding approach based on ITS2 biomarker.

For DHW preparation, we detected 35 PHS covering 25 PHMs, including the processed herbal materials such as Chao Baishu, the sensitivity of PHMs (define as the ratio of the detected prescribed herbal materials and the prescribed herbal materials in theory) was 69.4% based on ITS2 (**Table 4**), which was the largest number of detected PHS in this work. Among the detected PHS from 18 samples, 15 of the 35 detected PHS were found with an average relative abundance over 0.1%, where seven PHS were identified with an average relative abundance over 1%, including *Angelica sinensis* (2.0%), *Asarum sieboldii* (1.2%), *Notopterygium franchetii* (1.9%), *Notopterygium incisum* (1.8%), *Paeonia lactiflora* (5.3%), *Paeonia veitchii* (2.0%) and *Pogostemon cablin* (3.7%). Three PHS (*Clematis hexapetala, Coptis teeta, Paeonia lactiflora*) were found in all samples, but highly enriched in DHW.A samples. Average of the relative abundance of *Glycyrrhiza uralensis* (1.56%) and *Osmunda japonica* (1.64%) detected in samples from DHW.A was 1.6 times more than DHW.B samples (the relative abundance of these species in DHW.B was 0.94% and 0.98%, respectively), while *Coptis deltoidei* (one reads in DHW.A.III3), *Ephedra intermedia* (three reads in DHW.A.III2), *Gastrodia elata* (one reads in DHW.A.III3) and *Rheum tanguticum* (three reads in DHW.B.III1) were only detected from one sample. Noticeably, the substituted herbal species *Anemone nemorosa* (0.31%) with the same genus with PHS, was found with high relative abundance in most samples, especially in DHW.A.II and DHW.A.III.

**Table 4.**
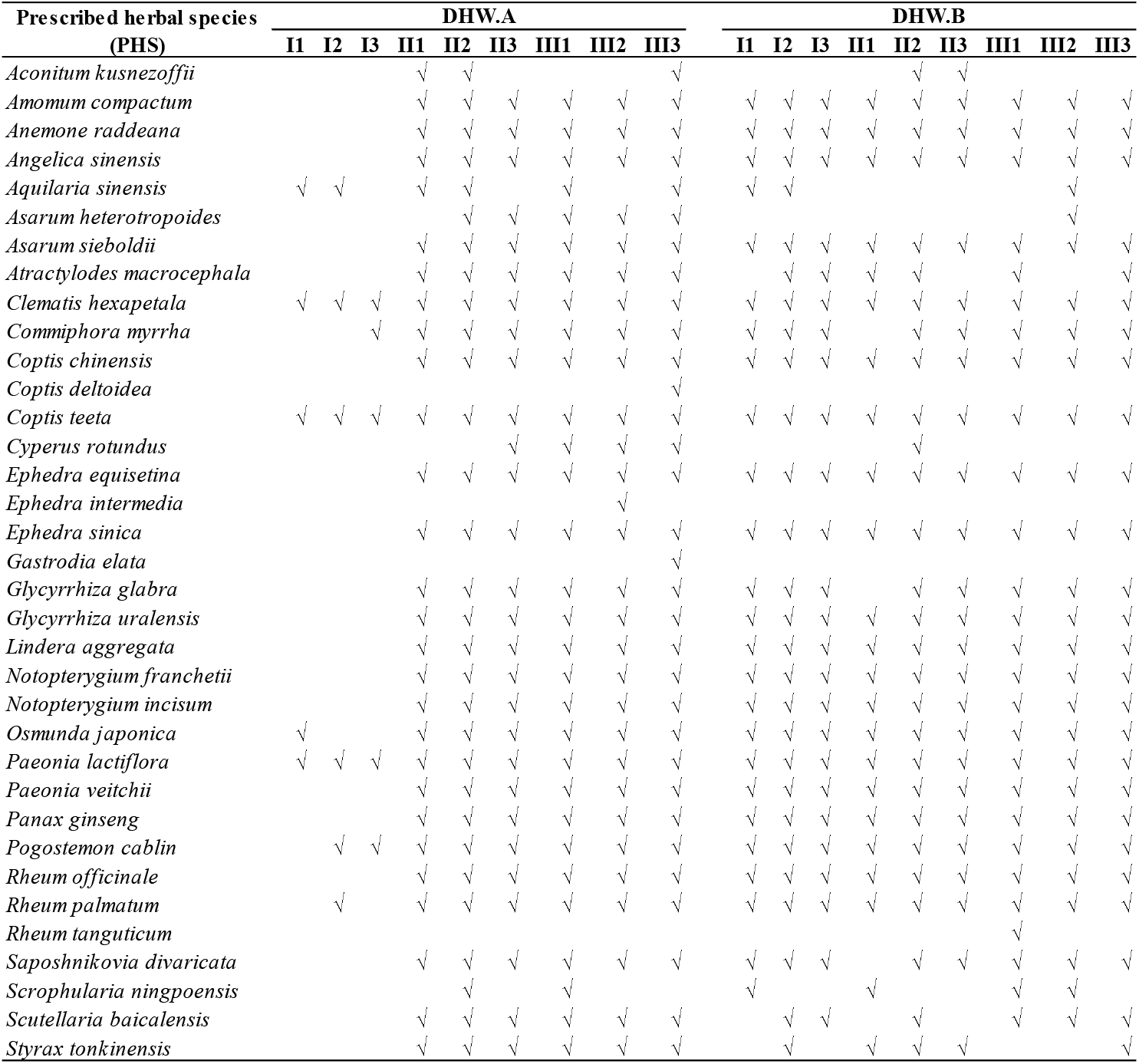
Prescribed herbal species for DHW preparation and their presence in each sample by multi-barcoding approach based on ITS2 biomarker.

#### Analysis of herbal materials in the TCM preparations based on *trnL*

For NJW, seven PHS belonged to four genera were detected with low abundance, including *Codonopsis pilosula, Curcuma kwangsiensis*, *Curcuma longa, Curcuma phaeocaulis, Nardostachys chinensis, Nardostachys jatamansi, Scutellaria baicalensis* (**Table 5**), among them, *Nardostachys chinensis* was captured in all samples, while *Codonopsis pilosula* and *Nardostachys jatamansi* were only identified in one sample with one reads, which suggested that DNA of these low relative abundance species was hard to be extracted or the *trnL* c/h primers were not suitable enough for the determination of *Codonopsis pilosula* and *Nardostachys jatamansi*. The substituted *Astragalus* (3.9%) and *Mentha* (8.1%) were captured with high relative abundance. As for possible contaminated herbal species, they were dispersedly distributed in 52 genera.

**Table 5.** Prescribed herbal species for NJW preparation and their presence in each sample by multi-barcoding approach based on *trnL* biomarker.

| Prescribed herbal species<br>(PHS) | NJW.A |  |  |  |  |  |  |  |  | NJW.B |  |  |  |  |  |  |  |
| --- | --- | --- | --- | --- | --- | --- | --- | --- | --- | --- | --- | --- | --- | --- | --- | --- | --- |
|  | I1 | I2 | I3 | II1 | II2 | II3 | III1 | III2 | III3 | I1 | I2 | I3 | II1 | II2 | II3 | III1 | III3 |
| *Nardostachys chinensis* | √ | √ | √ | √ | √ | √ | √ | √ | √ | √ | √ | √ | √ | √ | √ | √ | √ |
| *Nardostachys jatamansi* |  |  |  |  |  |  |  | √ |  |  |  |  |  |  |  |  |  |
| *Scutellaria baicalensis* |  |  |  |  |  |  |  |  | √ | √ |  | √ | √ |  | √ |  | √ |
| *Curcuma kwangsiensis* | √ |  |  | √ |  |  |  |  |  | √ |  |  |  |  |  | √ |  |
| *Curcuma longa* | √ |  |  | √ | √ |  |  |  |  | √ | √ | √ |  |  |  | √ |  |
| *Curcuma phaeocaulis* | √ |  |  | √ | √ |  |  | √ | √ | √ | √ | √ |  |  |  | √ |  |
| *Codonopsis pilosula* |  |  |  |  |  |  |  |  |  |  |  | √ |  |  |  |  |  |

Because of complex biological ingredients of DHW, the sensitivity of PHMs (18 PHMs, 22 PHS) was only 50% based on *trnL*. Among 22 detected PHS, 12 (**Table 6**) were detected in all samples with an average relative abundance greater than 0.1%, except *Coptis chinensis* (0.05%), in which six of them exceeded 1%, 10 of 22 PHS were below 0.05%. Moreover, the relative abundance of 22 PHS detected from DHW.A was higher than DHW.B. *Boswellia neglecta* (6.4%) was the dominate species, followed by *Glycyrrhiza uralensis* (4.2%), and then *Coptis deltoidei* (2.6%). Nevertheless, *Ephedra equisetina* (12 reads in DHW.A.II3 and 8 reads in DHW.A.III3) and *Scrophularia ningpoensis* (only one reads in both DHW.A.I2 and DHW.A.III1) were only found in two samples. The reason for this low abundant PHS might be due to the manufacturing process: several herbal materials (such as Chao Baishu, vinegar-process Xiangfu) need to be boiled or fried before adding into a TCM preparation.

**Table 6.**
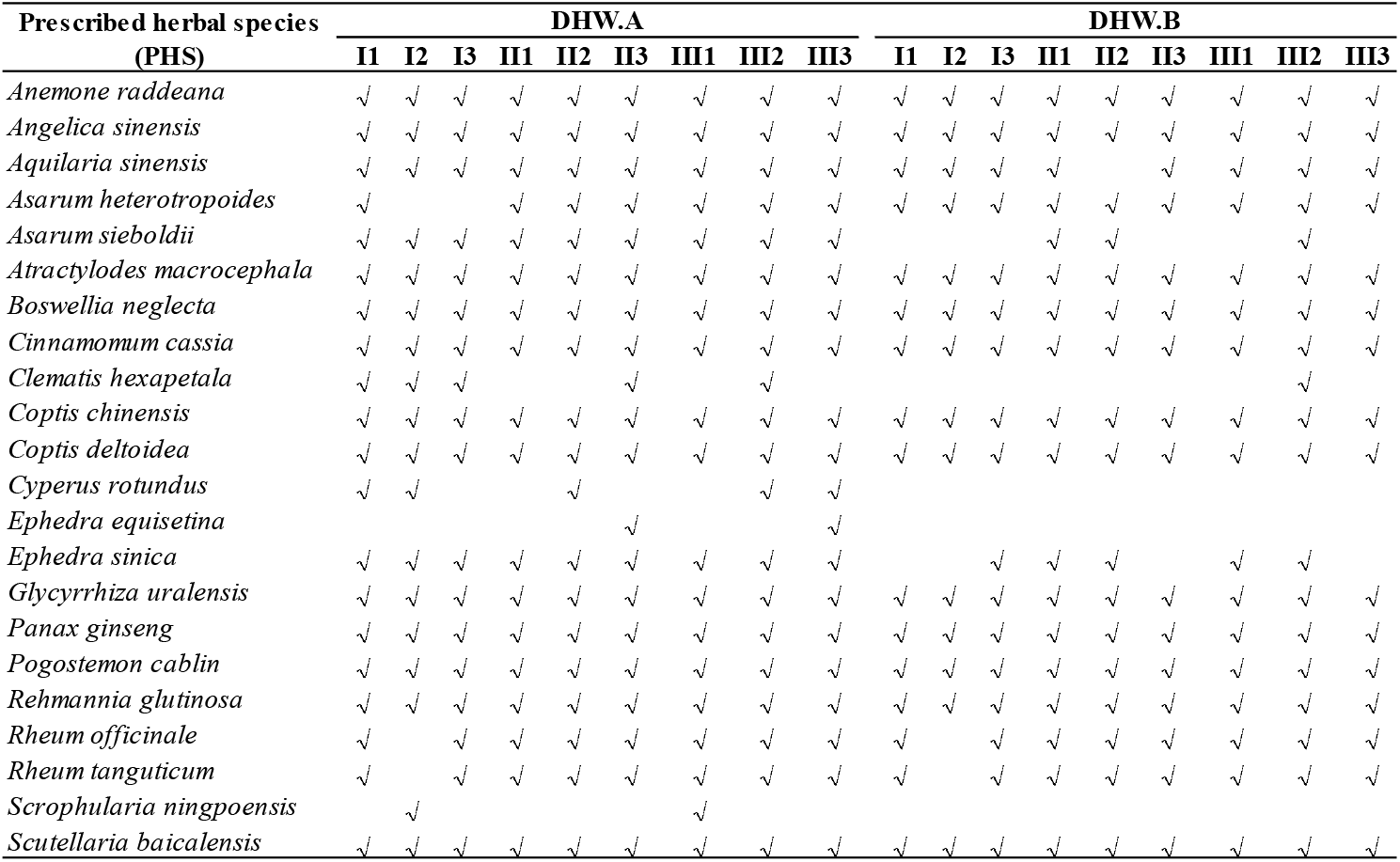
Prescribed herbal species for DHW preparation and their presence in each sample by multi-barcoding approach based on *trnL* biomarker.

The analysis of the sensitivity on BYW and YGW samples based on ITS2 and *trnL* biomarker was shown in **Supplementary Table 7-10**. Comparing the analysis result of DHW and NJW, NJW only contains one preprocessed PHM (the extractive of Huangqin), while DHW has seven preprocessed PHMs. This might be due to the more complex preprocessing procedure of DHW. Comparing the detecting result with ITS2 biomarker, much fewer species were identified using *trnL* biomarker, which might be due to DNA extraction, primer specification and the limitation of *trnL* database of Genbank. The three biological replicates from these batches have shown different prescribed herbal species (PHS) compositions based on both ITS2 or for *trnL* (**Table 3–6** and **Supplementary Table 7-10**), which might be potentially caused by DNA extraction, PCR amplification, high-through sequencing technology, and the previous researches of LDW^14^, YMW^15^, LXW^16^ and JQW^17^ have also shown this phenomenon.

All detected species including PHS, SHS and CHS of these four TCM preparations were also provided in **Supplementary Table 11** (provided as a separate attachment in .xlsx format). Based on ITS2 biomarker, we detected 8, 25, 9 and 6 prescribed herbal materials of BYW, DHW, NJW and YGW, respectively. The detected proportion of prescribed herbal materials was 100% for BYW and NJW, followed by DHW (69.4%) and YGW (66.7%). As for *trnL*, 5, 18, 4 and 4 prescribed herbal materials of BYW, DHW, NJW and YGW were respectively detected, and the maximum sensitivity of prescribed herbal materials was 62.5% among the four TCM preparations in this experiment. The analysis strongly suggested the multi-barcoding approach has a high sensitivity in identifying prescribed herbal materials of TCM preparations, especially based on ITS2 dataset.

### 3.4. Prediction model to predict the identity and quality of TCM preparations

By enabling a model to differentiate the sample from a different group, we can also identify the manufacturer, the batch from where samples were collected. Various distance measures can be used to evaluate the inter/intra-manufacturers difference. Here, we calculated the Euclidean distances of any two samples based on the existence of all detected species and then clustered the samples according to their similarity. We took DHW as a case study. The results showed that most samples from DHW.A and DHW.B clustered together respectively based on both ITS2 (**Figure 3A** and **B**) and *trnL* (**Figure 3C** and **D)** biomarkers, suggesting that the high similarity of intra-manufacturer samples. It is obvious that DHW.A.II and DHW.A.III is clustered with DHW.B samples, whereas three samples of DHW.A.I were gathered and distant from the other samples (**Figure 3A and B**). The reason for such separation might be the existence of substituted herbal species such as *Senna, Amaranthus, Glycine* and contaminated herbal species such as *Arachis, Brassica, Solanum* and *Oryza*. As for NJW (**Supplementary Figure 6**), the samples from two manufacturers (A&B) were scattered, based on either ITS2 or *trnL*, while clustered tighter within batches, which depicted the high consistency between batches of NJW samples. The cluster analysis of BYW and YGW samples was shown in **Supplementary Figure 7-8** respectively, which showed the clear difference between manufacturers, as well as high similarly within the same manufacturer.

**Figure 3.**
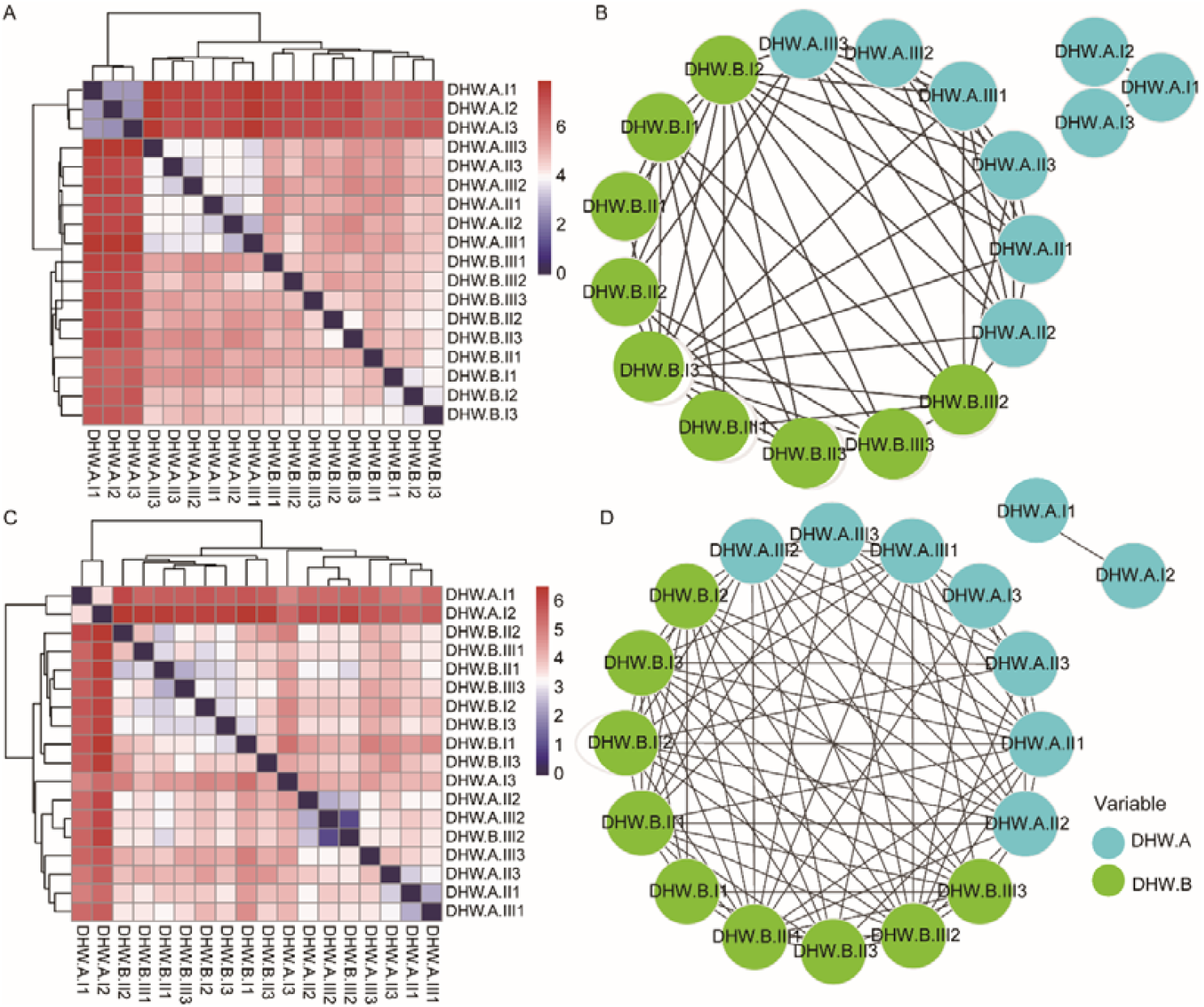
Comparison of the similarity of all DHW samples from intra-/inter-manufacturers based on prescribed herbal materials using Euclidean distances. Heatmap clusters displayed the distance of all samples based on the existence of prescribed herbal species using hierarchical clustering, and network clusters illustrated these differences in Cytoscape based on ITS2 (A and B) and *trnL* (C and D) sequencing results. For heatmap (A & C), the gradient color bars mean the distance between any two samples, while the red and the blue color depicts the two extreme distances between samples. For network (B & D), each edge represents the distance of any two samples with distance less than or equal to 5.0 for ITS2 and 4.2 for *trnL*.

PCA analysis was also performed to explore the consistency of samples from two manufacturers. The samples from DHW.B were clustered more closely than DHW.A based on ITS2 and *trnL* biomarker. Based on ITS2, the samples of DHW from intra-batch were clustered together, and the inter-batches distributed sparsely, whereas based on *trnL*, the samples of DHW.A were dispersed far apart (**Supplementary Figure 9C and D**), which suggested that the consistency of DHW.B samples was better than DHW.A. The cluster degree of samples from NJW (**Supplementary Figure 9E** and **F**) was more dispersive than DHW. The result of BYW and YGW (**Supplementary Figure 9 A&B** and **G&H**) was also showed similar results.

To explore which species drove the difference of intra-/inter-manufacturers samples, LEfSe analysis was conducted. 13 prescribed herbal species from DHW.A and four of DHW.B (**Figure 4A**) were identified as tentative biomarkers. Through mRMR, five PHS from DHW.A and two PHS of DHW.B were selected and visualized in ROC curves (**Figure 4B**) to evaluate their classification ability. As the curve of *Glycyrrhiza glabra* was below the model score curve, we removed this biomarker from DHW.A. Thus, *Coptis chinensis, Ephedra equisetina, Lindera aggregate* and *Panax ginseng* were chosen as unique biomarkers of DHW.A, whereas *Rheum palmatum* and *Clematis hexapetala* were selected as representative biomarkers of DHW.B. All of them are of high discrimination power, which could be used separately or in combination to differentiate the samples from the two manufacturers.

**Figure 4.**
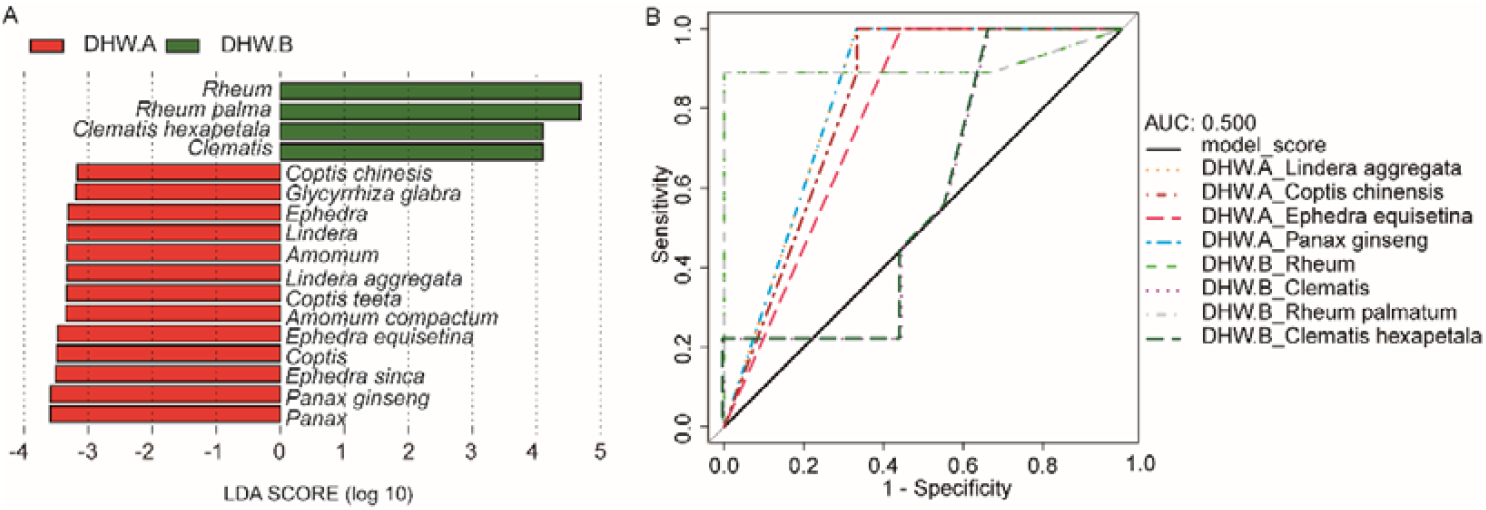
The difference of samples from the two manufacturers (A & B) could be driven by a few discriminative prescribed herbal species of prescribed herbal materials of DHW using ITS2 biomarker. (A) The legacy biomarkers selected by LEfSe; (B) ROC curves to evaluate the effect of the legacy biomarkers after removing redundant markers from the two manufacturers.

### 3.5. Comparison of ITS2 and *trnL* on resolutions and sensitivities

Through detecting their prescribed herbal species, the detected proportion of prescribed herbal materials was 100% for BYW and NJW, followed by DHW (69.4%) and YGW (66.7%) based on ITS2, while 62.5%, 50%, 44.4% and 44.4% for BYW, DHW, NJW and YGW based on *trnL* datasets respectively (**Table 7**). The sensitivities of ITS2 is better than *trnL* in all TCM preparations, but *trnL* biomarker could also detect the PHS of PHMs that ITS2 couldn’t, and the union of both increases the sensitivity of the lower limit to 77.8%, providing a more reliable (as for positive detections) detected result.

**Table 7.** The sensitivity of prescribed herbal materials for four TCM preparations based on ITS2 and *trnL* biomarker.

|  | ITS2 (%) | *trnL* (%) | Union (%) |
| --- | --- | --- | --- |
| BYW | 100 | 62.5 | 100 |
| DHW | 69.4 | 50 | 77.8 |
| NJW | 100 | 44.4 | 100 |
| YGW | 66.7 | 44.4 | 77.8 |

Note that the sensitivity was defined as the ratio of the detected prescribed herbal materials and the prescribed herbal materials could be detected in theory.

As can be observed from the Venn diagram (**Figure 5**), the detection result of BYW, all its prescribed herbal materials were detected. As for DHW, the union detection result of these two regions was 38 PHS, covering 28 prescribed herbal materials, which increased the identification efficiency to 77.8%. Similarly, the detection result of *trnL* from NJW preparation was a subset of ITS2, with 100% sensitivity. For YGW samples, the union of these two regions increased the sensitivity to 77.8%, because of two undetected PHMs named Myristicae semen (Rougui) and Dioscoreae rhizome (Shanyao). This result has also confirmed the high reliability of the multi-barcoding approach. We then compared our result with the previous studies, including JQW, LXW, YMW and the YYW (**Table 8**), which indicated the reliability of the multi-barcoding approach, this was also suggested that the complexity of biological ingredients of TCM preparation has also negatively affected the detected results.

**Figure 5.**
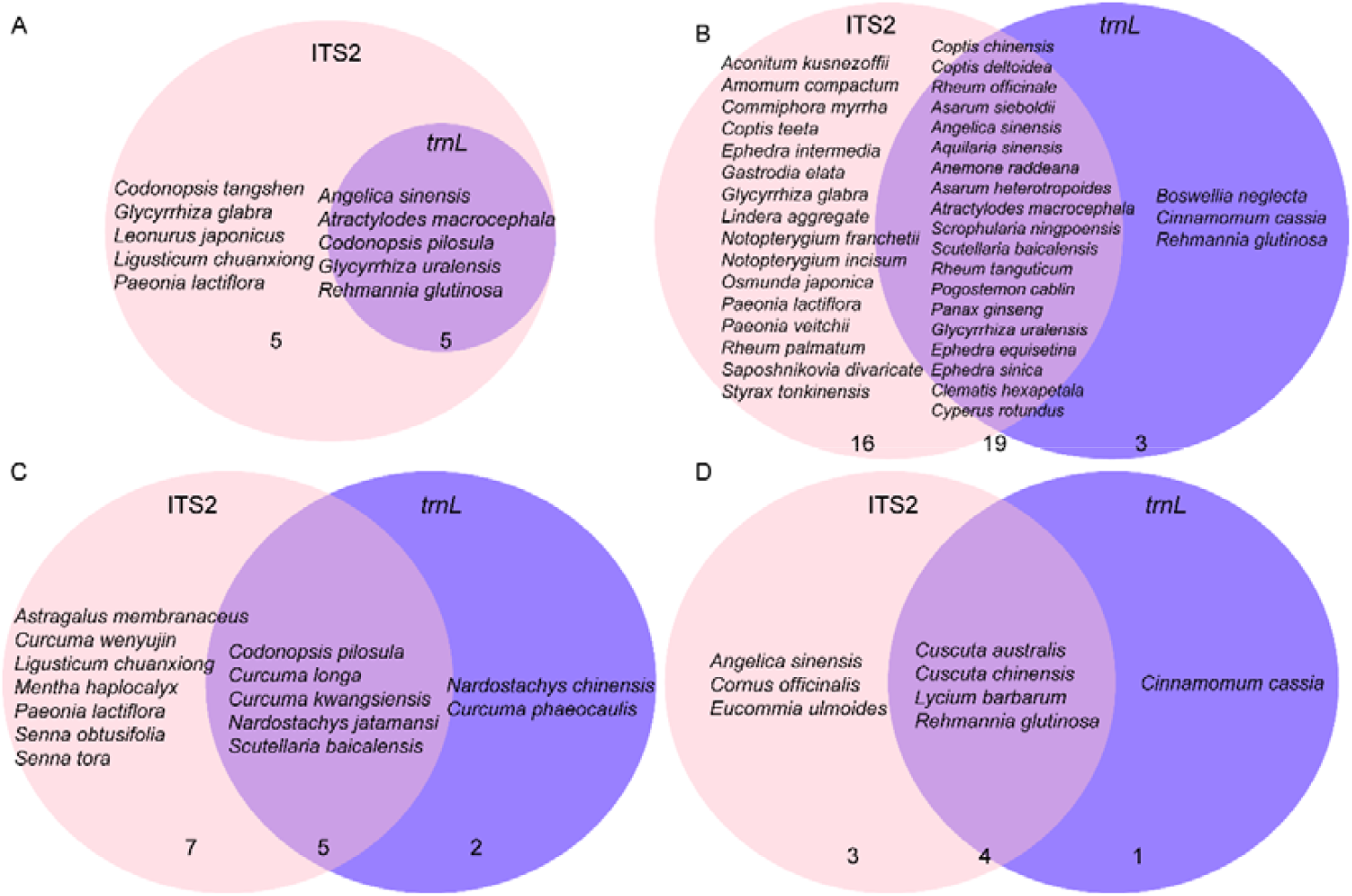
The identified specific and shared prescribed herbal species of TCM preparations based on ITS2 and *trnL*. Results on (A) BYW; (B) DHW; (C) NJW; (D) YGW were shown. The numbers below the Venn diagram mean the number of prescribed herbal species detected based on ITS2 only, *trnL* only and the intersection of the two.

**Table 8.**
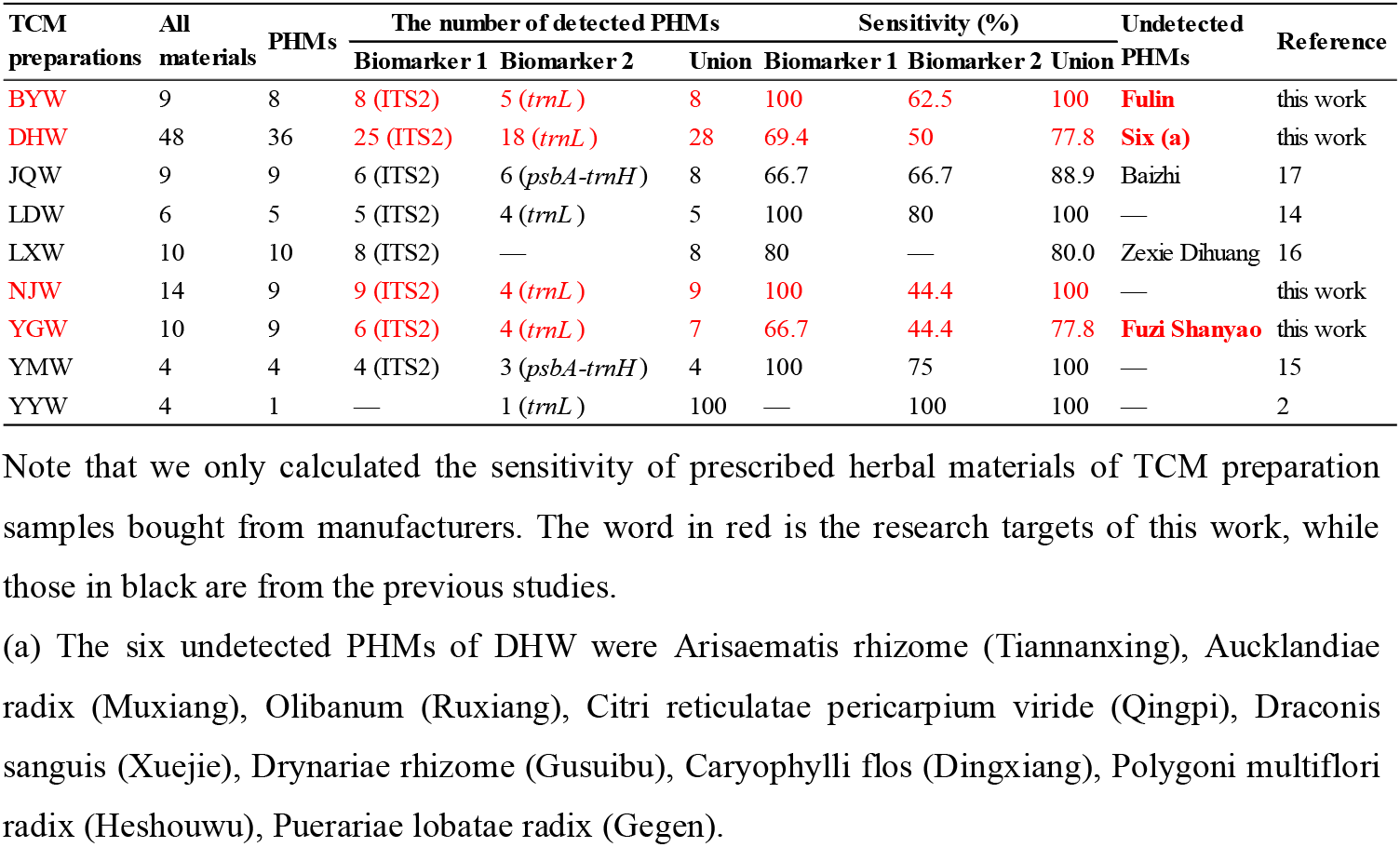
Comparison of the sensitivity of prescribed herbal materials through detected prescribed herbal species of TCM preparations.

| TCM preparations | All materials | PHMs | The number of detected PHMs |  |  | Sensitivity (%) |  |  | Undetected PHMs | Reference |
| --- | --- | --- | --- | --- | --- | --- | --- | --- | --- | --- |
|  |  |  | Biomarker 1 | Biomarker 2 | Union | Biomarker 1 | Biomarker 2 | Union |  |  |
| BYW | 9 | 8 | 8 (ITS2) | 5 (*trnL*) | 8 | 100 | 62.5 | 100 | **Fulin** | this work |
| DHW | 48 | 36 | 25 (ITS2) | 18 (*trnL*) | 28 | 69.4 | 50 | 77.8 | **Six (a)** | this work |
| JQW | 9 | 9 | 6 (ITS2) | 6 (*psbA-trnH*) | 8 | 66.7 | 66.7 | 88.9 | Baizhi | 17 |
| LDW | 6 | 5 | 5 (ITS2) | 4 (*trnL*) | 5 | 100 | 80 | 100 | — | 14 |
| LXW | 10 | 10 | 8 (ITS2) | — | 8 | 80 | — | 80.0 | Zexie Dihuang | 16 |
| NJW | 14 | 9 | 9 (ITS2) | 4 (*trnL*) | 9 | 100 | 44.4 | 100 | — | this work |
| YGW | 10 | 9 | 6 (ITS2) | 4 (*trnL*) | 7 | 66.7 | 44.4 | 77.8 | **Fuzi Shanyao** | this work |
| YMW | 4 | 4 | 4 (ITS2) | 3 (*psbA-trnH*) | 4 | 100 | 75 | 100 | — | 15 |
| YYW | 4 | 1 | — | 1 (*trnL*) | 100 | — | 100 | 100 | — | 2 |

Note that we only calculated the sensitivity of prescribed herbal materials of TCM preparation samples bought from manufacturers. The word in red is the research targets of this work, while those in black are from the previous studies.
(a) The six undetected PHMs of DHW were *Arisaematis rhizome* (Tiannanxing), *Aucklandiae radix* (Muxiang), *Olibanum* (Ruxiang), *Citri reticulatae pericarpium viride* (Qingpi), *Draconis sanguis* (Xuejie), *Drynariae rhizome* (Gusuibu), *Caryophylli flos* (Dingxiang), *Polygoni multiflori radix* (Heshouwu), *Puerariae lobatae radix* (Gegen).

Though the sensitivity and reliability of multi-barcoding approach have been clearly demonstrated, the results on ITS2 and *trnL* are clearly different. It is obvious that ITS2 showed a higher sensitivity than that of *trnL* for PHMs detection. The reason might be due to the longer conserved region of ITS2 which can capture more information. Nevertheless, the role of *trnL* is irreplaceable, as it could complement ITS2 for more reliable identification of the prescribed herbal materials of TCM preparations such as for the biological ingredient analysis of DHW and YGW in this work.

## 4. Discussions and Conclusion

As already know to us, herbal materials are the most important elements in different traditional medicines. An increasing number of papers on DNA-based authentication of single herbs have been published^27,39–44^, while a few applications of the multi-barcoding approach for TCM preparations were reported^14,45–47^.

### 4.1. The multi-barcoding approach decoded the prescribed herbal materials of the four TCM preparations

In this work, we have systematically examined the universality, sensitivity and reliability of multi-barcoding approach for four representative TCM preparations. This method has successfully detected the species (including prescribed, substituted and contaminated species) contained in a sample with high sensitivity, indicating the good universality of the method and its potential value for daily TCM supervision. As we could determine the existence of all species contained in one sample at species level, these results have indicated an adequate sensitivity of this method in decoding herbal materials of TCM preparations through authenticating their corresponding species. The combined results of ITS2 and *trnL* have increased the sensitivity from 77.8% to 100% that highlights the practical application value and high reliability of this approach. Particularly, the ITS2 exhibited an excellent ability and sensitivity for identifying herbal materials. Although the resolution of *trnL* was lower than that of ITS2, it could also be used to reinforce or complement ITS2 for more reliable results. These results have demonstrated that multi-barcoding was an efficient tool for decoding the herbal materials of various kinds of TCM preparations.

For example for BYW and NJW, all prescribed herbal materials were detected through authenticating its corresponding prescribed herbal species. The detected prescribed herbal species of DHW were 35 (covered 25 prescribed herbal materials), 22 (covered 18 prescribed herbal materials) based on ITS2 and *trnL*, respectively. The union dataset of ITS2 and *trnL* has boosted the sensitivity increasing from 69.4% to 77.8% for DHW samples. However, six prescribed herbal materials were not detected in all DHW samples based on either ITS2 or *trnL*. These phenomena might be due to various preprocessing procedures, such as decocted or stir-fried herbal materials, who’s DNA was damaged or degraded. We also note that due to several influencing factors, such as geological location, cultivation conditions, climate and other conditions, the sensitivity of PHMs of each TCM preparation sample is different.

This multi-barcoding approach has successfully analyzed the herbal materials of four TCM preparations, which could not be realized through traditional methods, such as morphological and biochemical means. In the future, more diverse sets of TCM preparations could be assessed by this method, which not only making the identification of TCM preparation automatically, but also accelerating the digitization and modernization of TCM management process.

### 4.2. Outlook and future plans

However, a deeper and more comprehensive improvement of this multi-barcoding approach still needs to be carried out. A more comprehensive species database was necessary, since the reliability of the biological ingredient analysis method for TCM preparation were largely dependent on the coverage of the reference database^2^. In our future study, we can utilize multiple databases, including the GenBank database, as well as tcmbarcode database^48^, EMBL, DDBJ and PDB^2^ to obtain more complete results. Additionally, more biomarker candidates can be considered for assessing the quality of TCM preparation.

Firstly, the multi-barcoding approach could be an attempt to use in identifying the animal materials, because the animal materials still are an important component of TCM, are often combined with medical herbs to exert its pharmacological effects^49^.

Secondly, chemical ingredients and biological ingredients are indivisible yet both important for quality assessments of TCM preparations. Therefore, combining the chemical methods with DNA barcoding approach, the detection of TCM ingredients will outperform than the results of any one of them. Although this thought was initially tested by our group^11^, there is still room for further improvement.

Thirdly, the network pharmacology approach has provided us with a more direct view about the drug-target interactions^50^, which gives us an insight into how to optimize the existing drugs and to discover the new medicine for satisfying the requirements of overcoming complex diseases. Thus, the pharmacological usage should be considered in the QC of TCM preparations, especially for the specific usages of TCM, such as the mechanism-based QC of YIV-906^51^. This theory has also inspired us to explore the potential treatments of COVID-19 from biological ingredients of TCM preparations^52^. In fact, the ingredients such as Glycyrrhizae Radix Et Rhizoma could frequently interact with the target of COVID-19: ACE2^20,52^. Through data-mining, the characteristic of eight biological ingredients of DHW is corresponding to the classic Warm disease’s symptoms of syndrome differentiation of COVID-19, which might prove effective for the treatment of COVID-19^52^. These biological ingredient information, if combine with public health data, might shed more lights on the susceptibility of patient who has taken these TCM preparations, especially those elderly people.

Finally, many herbal medicines are taken orally^53^, undoubtedly exposed to the whole gastrointestinal tract microbiota, which provides sufficiently spatiotemporal opportunities for their direct or indirect interactions. For example, berberine, the major pharmacological ingredients of *Coptidis rhizome* (Huanlian)^54^, it promotes the production of short-chain fatty acid to shift the gut microbiota structure, while the poorly solubilized berberine^55^ was converted into dihydroberberine through a reduction reaction mediated by bacterial nitroreductase, then recovered to the original form after penetrating into the intestinal wall tissues^56^, through interactions, the microbial diversity in high-fat diet mice intestines was profoundly decreased^57^.

We believe that all of these efforts on QC of TCM preparations could and would joint-force and provide much better and optimized approaches for the next-generation TCM preparation quality control system. Through reshaping the symbiotic microbiome composition, we could provide novel therapeutic strategies to accelerate the realization of personalized therapeutics.

## Supporting information

9 pictures and 10 tables

## Acknowledgments

This work was partially supported by National Science Foundation of China grant 81573702, 81774008, 31871334 and 31671374, and National Key Research and Development Program of China grant 2018YFC0910502.

## Authors’ contributions

KN and HB designed the whole study. HB, MZH, CYC and QY collected the samples and conduced the DNA extraction and sequencing. XZ analyzed the sequencing data. XZ, HB, MZH and KN wrote, revised and proof-read the manuscript. All authors read and approved the final manuscript.

## Competing financial interests

The authors declare no competing financial interest.

## Data availability

The raw sequencing data used in this work was deposited to NCBI SRA database with accession number PRJNA562480. The ITS2 sequences of the sequenced single herbs were also deposited to NCBI SRA database with NCBI SRA database with accession number PRJNA600815.

## Notes

### Competing Interest Statement

The authors have declared no competing interest.

