## Supplementary material for "Decoding herbal materials of representative TCM preparations with the multi-barcoding approach": 9 pictures and 10 tables

**This file includes:**

Supplementary Figure 1-9

Supplementary Tables 1-10 (these nine Tables was shown in both Figure and Excel format)

Supplementary Tables 11 (only in .xlsx format)

References

**Supplementary Figures**

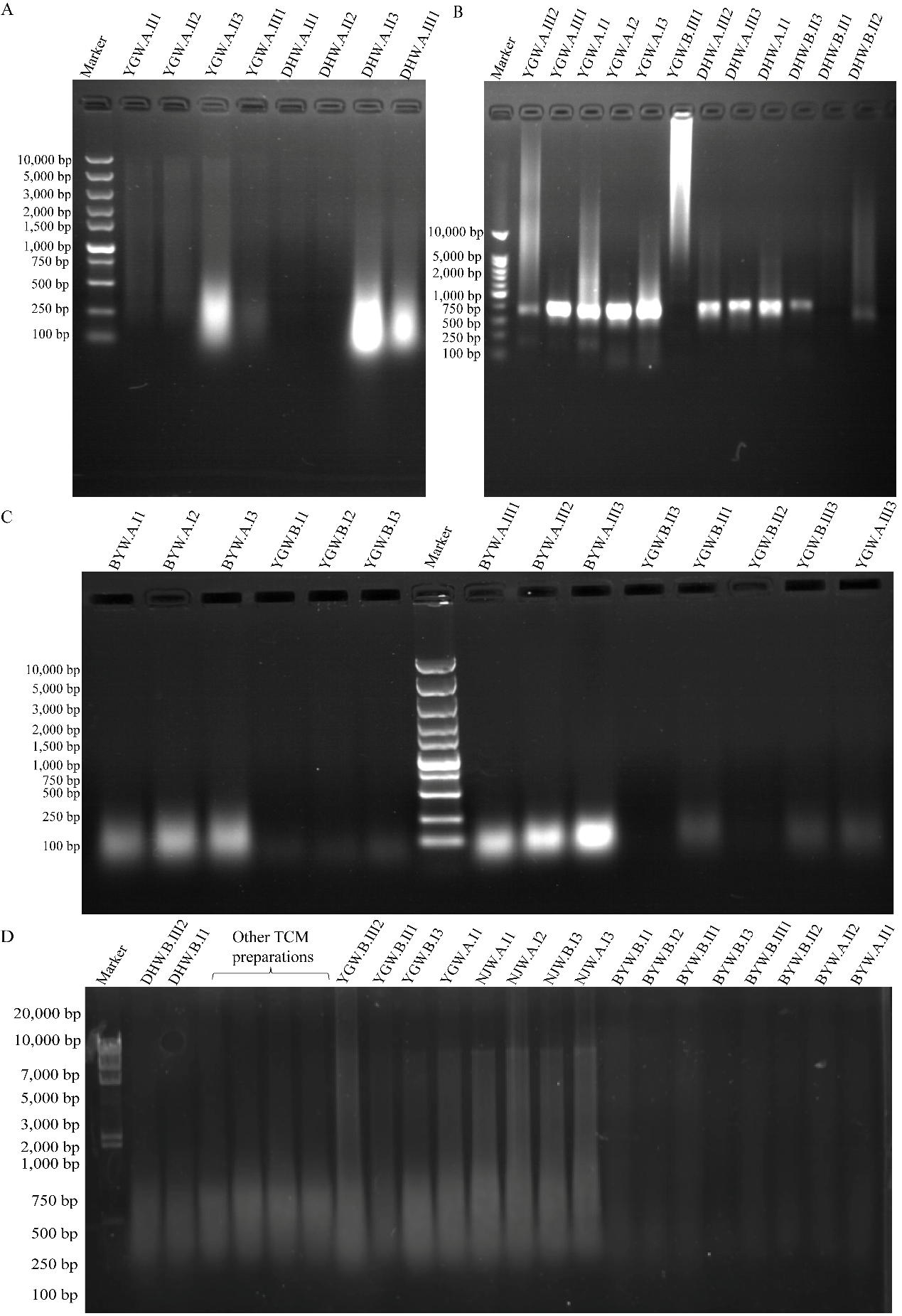

**Supplementary Figure 1. Genomic DNA extraction from TCM preparation samples by optimized cetyl trimethyl ammonium bromide (CTAB) method (TCM-CTAB).** (A) & (C) The DNA of Bazhen Yimu Wan (BYW) and Yougui Wan (YGW); (B) The DNA of Da Huoluo Wan (DHW) and Yougui Wan (YGW); (D) The DNA of Bazhen Yimu Wan (BYW), Da Huoluo Wan (DHW) and Yougui Wan (YGW), ‘Others’ mean the DNA of other TCM preparation.

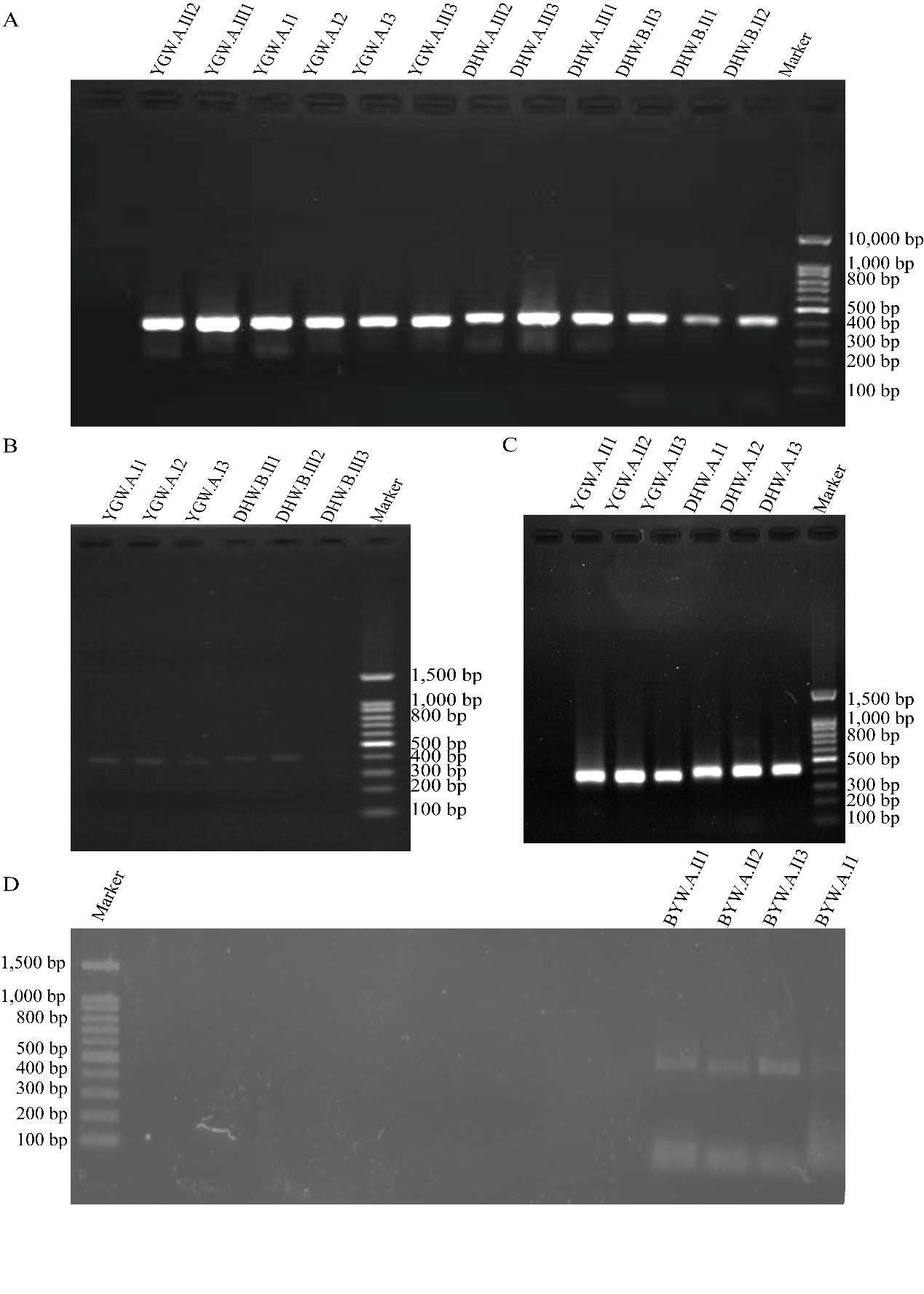

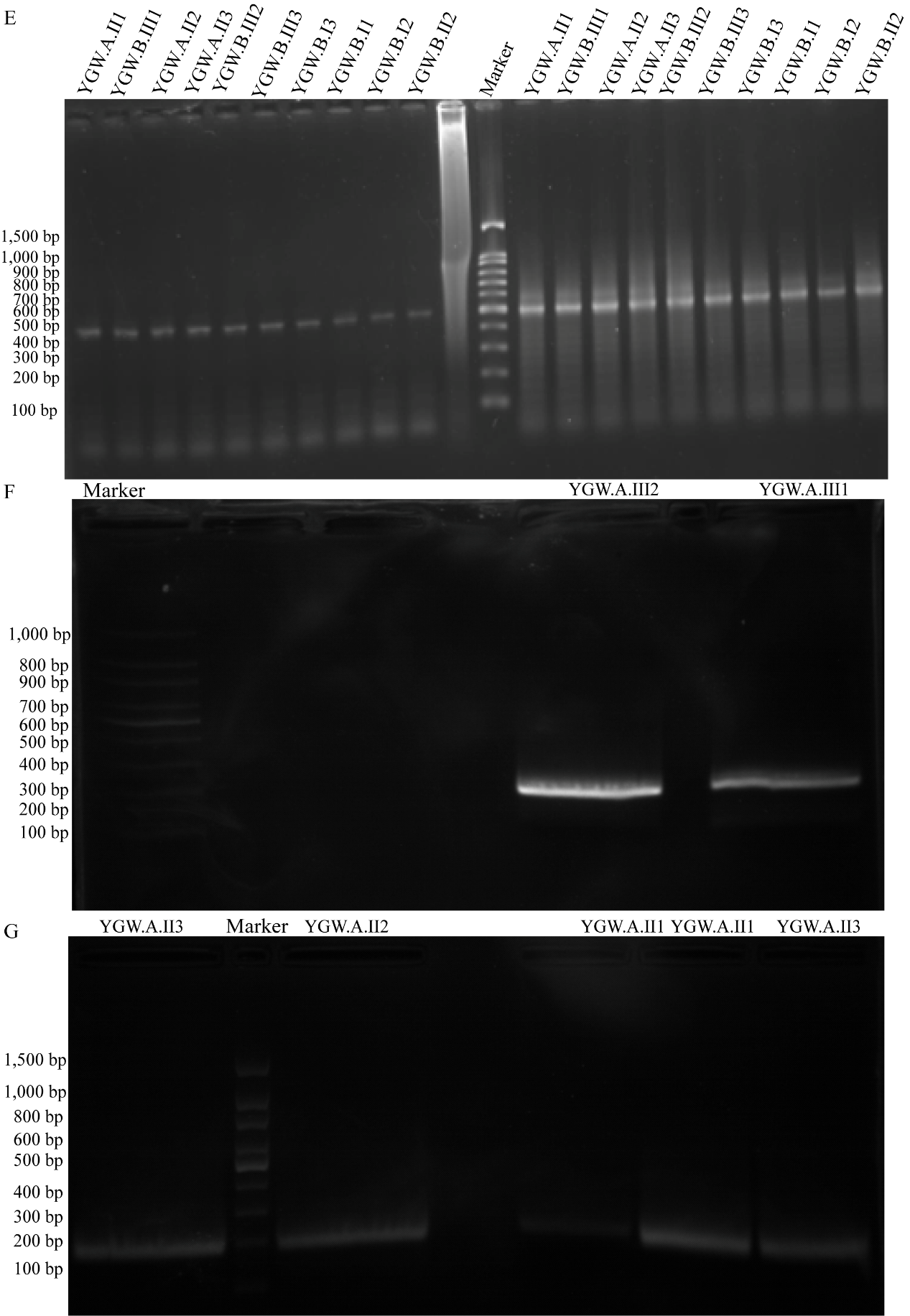

**Supplementary Figure 2. PCR amplification of TCM preparations.** (A) & (B) & (C) are the PCR amplification of Yougui Wan (YGW) samples and Da Huoluo (DHW) samples based on ITS2 biomarker, respectively; (D) is the PCR amplification of four Bazhen Yimu Wan (BYW) samples based on ITS2 biomarker; (E) is the PCR amplification of 10 YGW samples based on ITS2 biomarker at 1X buffer and 2X buffer, respectively; (F) & (G) are the PCR amplification of YGW samples based on *trnL* biomarker.

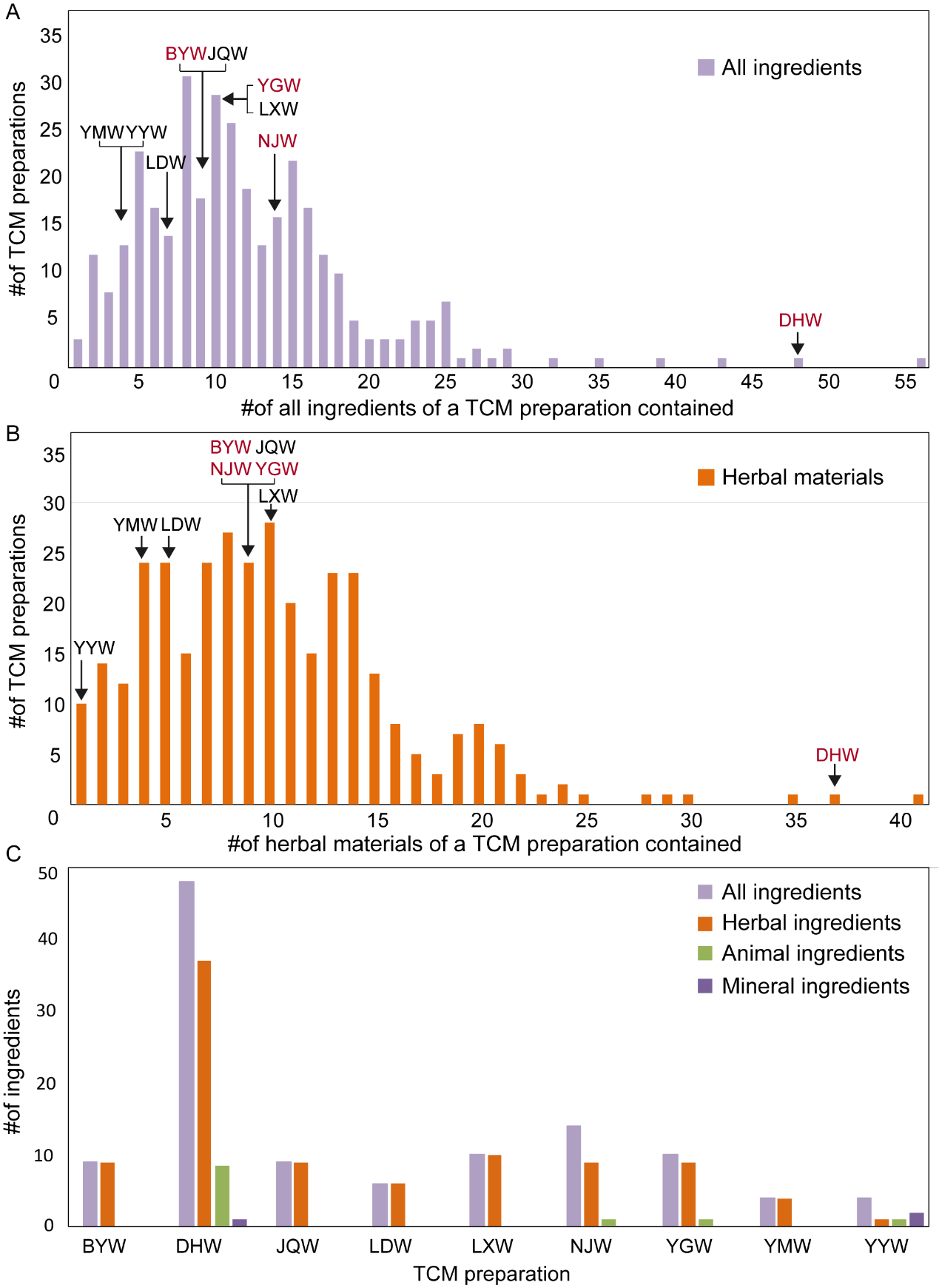

**Supplementary Figure 3. The distribution of all ingredients and the herbal materials only of all TCM preparations listed in Chinese pharmacopoeia.** (A) All ingredients of TCM preparation, including herbal, animal and mineral materials. (B) The herbal materials of a TCM preparation contained. The x-axis represents the number of all/herbal ingredients of a TCM preparation contained, y-axis means the corresponding number of the TCM preparations. The abbreviations are shown, from left to right, Yatong Yili Wan (YYW), Yimu Wan (YMW), Liuwei Dihuang Wan (LDW), Bazhen Yimu Wan (BYW), Yougui Wan (YGW), Niuhuang Jiangya Wan (NJW), Jiuwei Qianghuo Wan (JQW), Longdan Xiegan Wan (LXW) and Da Huoluo Wan (DHW), respectively. (C) The distribution of all ingredients and the detailed ingredient of nine TCM preparations. The abbreviations are shown, from left to right, Bazhen Yimu Wan (BYW), Da Huoluo Wan (DHW), Jiuwei Qianghuo Wan (JQW), Liuwei Dihuang Wan (LDW), Longdan Xiegan Wan (LXW), Niuhuang Jiangya Wan (NJW), Yougui Wan (YGW), Yimu Wan (YMW) and Yatong Yili Wan (YYW), respectively. The words marked in black are those already reported in previous studies, while the red is the research preparations used in this work.

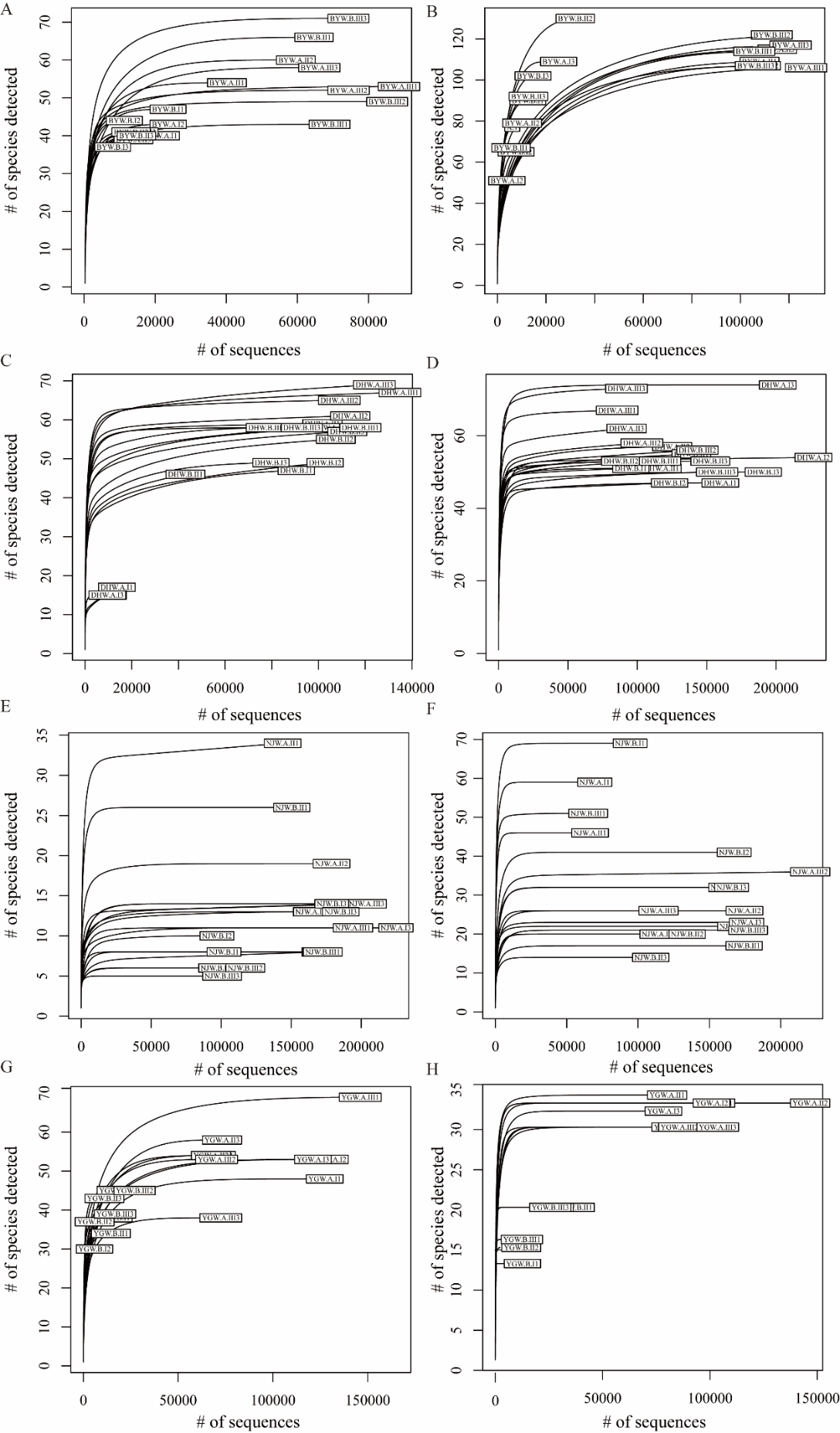

**Supplementary Figure 4. Rarefaction carves of sequencing depth sampled from two manufactures for per TCM preparation based on ITS2 and *trnL*.** (A) BYW based on ITS2; (B) BYW based on *trnL*; (C) DHW based on ITS2; (D) DHW based on *trnL*. (E) NJW based on ITS2. (F) NJW based on trnL; (G) YGW based on ITS2; (H) YGW based on *trnL*. The result shows all samples’ sequencing depth were saturation.

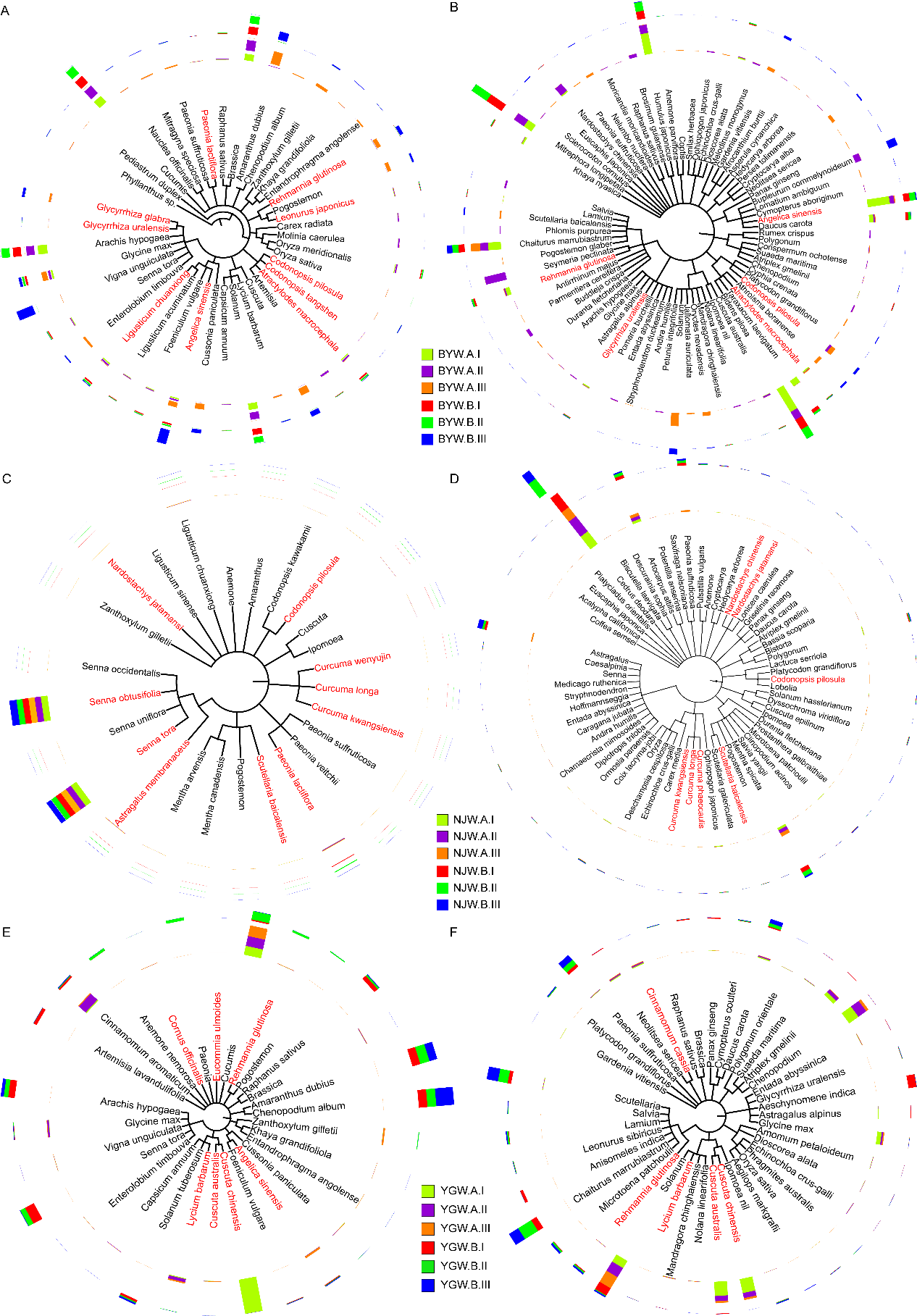

**Supplementary Figure 5. Phylogenetic analysis of the representative species that had at least 0.1% relative abundance for TCM preparation samples.** (A) BYW samples based on ITS2; (B) BYW samples based on *trnL*; (C) NJW samples based on ITS2; (D) NJW samples based on *trnL*; (E) YGW samples based on ITS2; (F) YGW samples based on *trnL*. The word marked in red means the prescribed herbal species, and the colorful bar means the average relative abundances of species across the three batches from the two manufacturers (A&B).

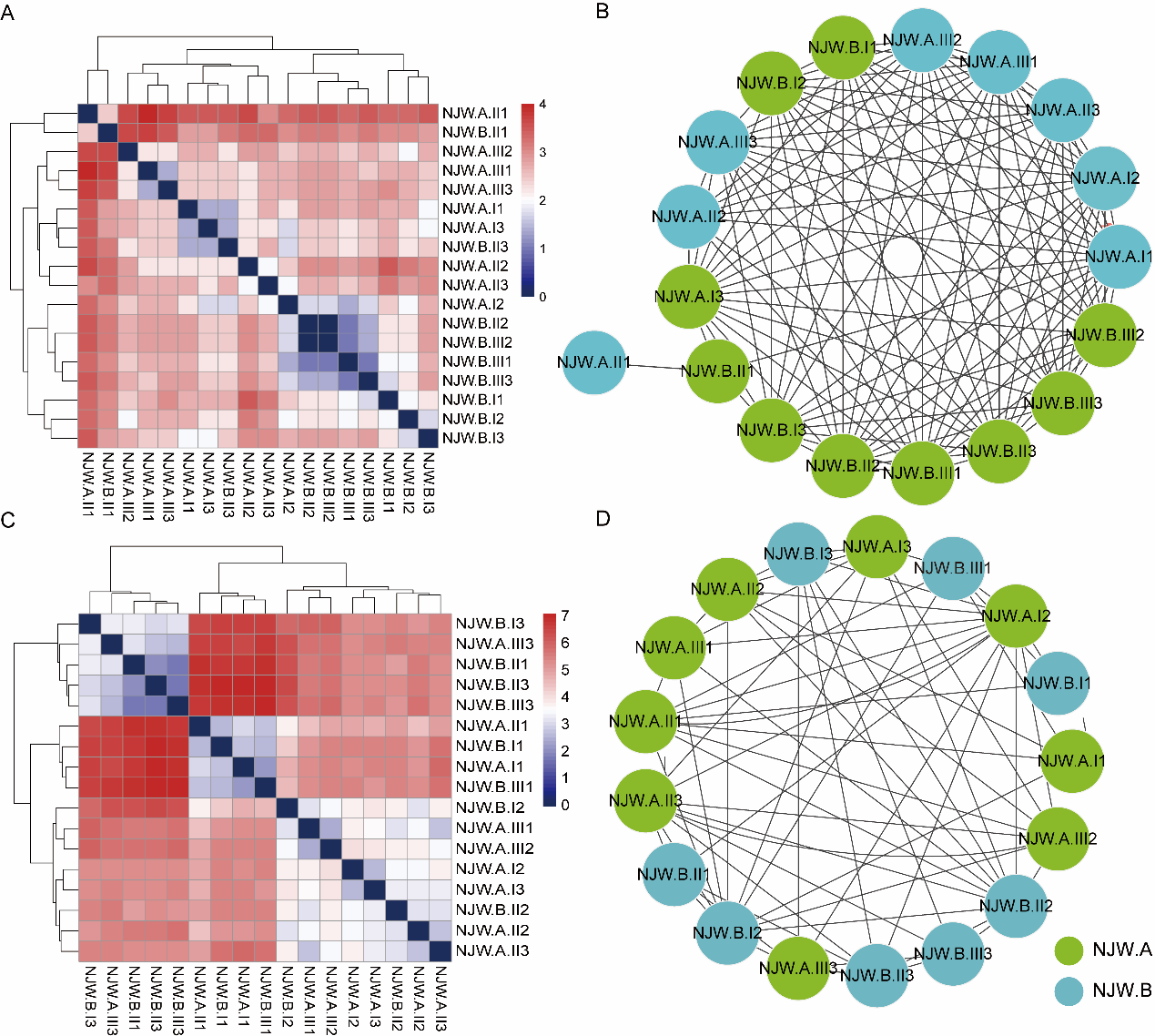

**Supplementary Figure 6. Comparison of the similarity of all NJW samples from intra-/inter-manufacturers based on prescribed herbal materials using Euclidean distances.** Heatmap clusters displayed the distance of all samples based on the existence of prescribed herbal species using hierarchical clustering, and network clusters illustrated these differences in Cytoscape based on ITS2 (A and B) and *trnL* (C and D) sequencing results. For heatmap (A & C), the gradient color bars mean the distance between any two samples, while the red and the blue color depicts the two extreme distances between samples. For network (B & D), each edge represents the distance of any two samples with distance less than 2.9 for ITS2 and 3.9 for *trnL*.

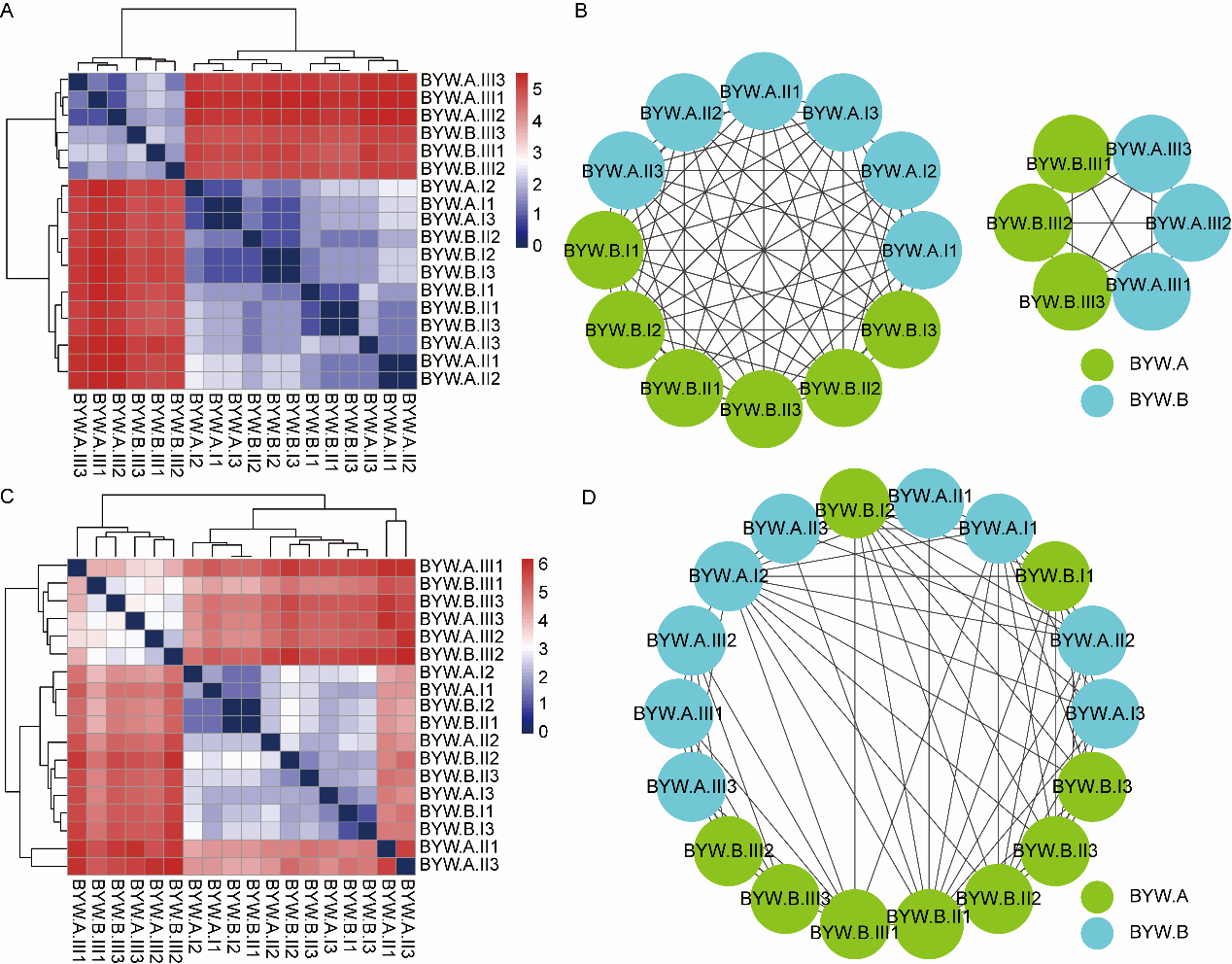

**Supplementary Figure 7. Comparison of the similarity of all BYW samples from intra-/inter-manufacturers based on prescribed herbal materials using Euclidean distances.** Heatmap clusters displayed the distance of all samples based on the existence of prescribed herbal species using hierarchical clustering, and network clusters illustrated these differences in Cytoscape based on ITS2 (A and B) and *trnL* (C and D) sequencing results. For heatmap (A & C), the gradient color bars mean the distance between any two samples, while the red and the blue color depicts the two extreme distances between samples. For network (B & D), each edge represents the distance of any two samples with distance less than 2.5 for ITS2 and 4.6 for *trnL*.

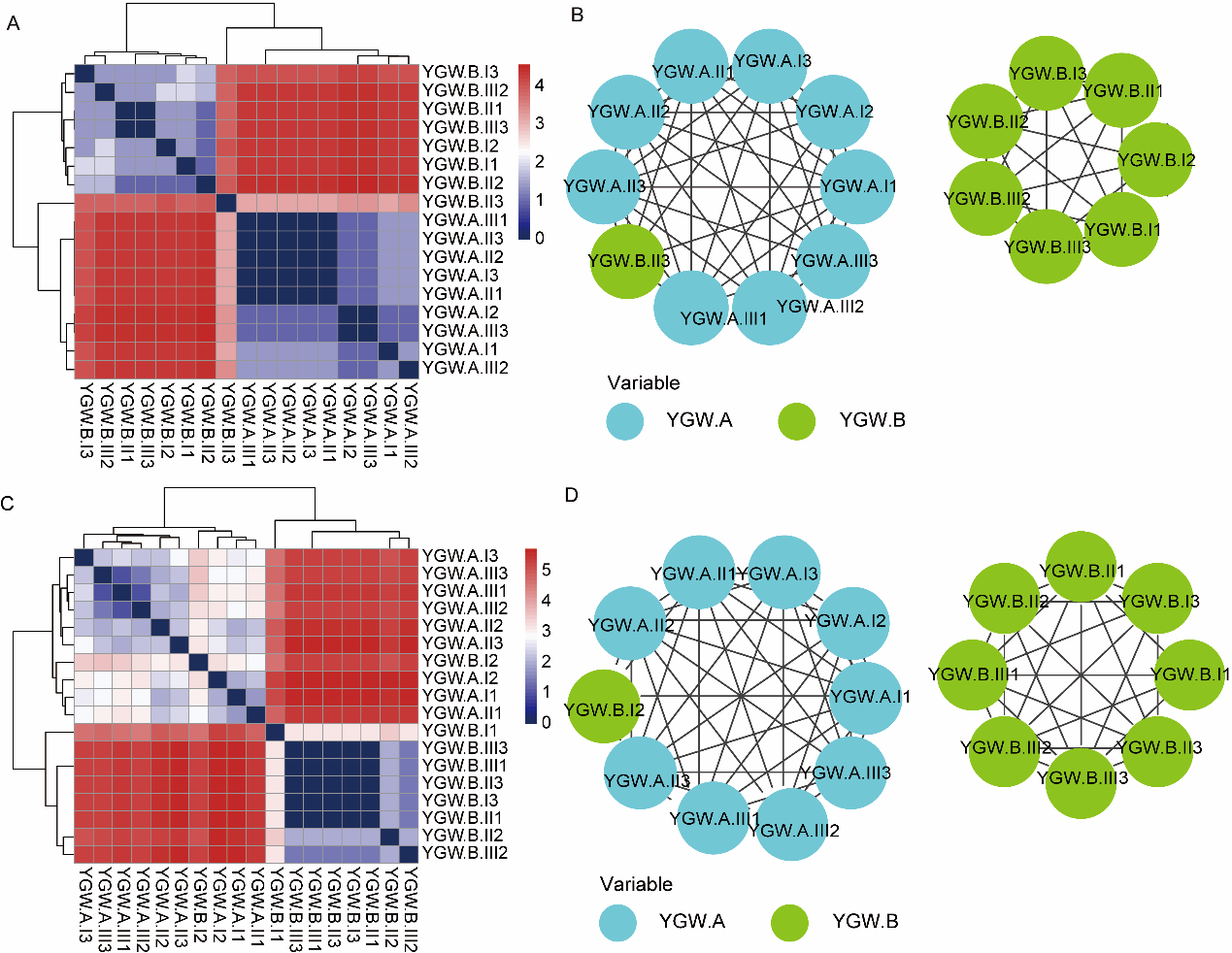

**Supplementary Figure 8. Comparison of the similarity of all YGW samples from intra-/inter-manufacturers based on prescribed herbal materials using Euclidean distances.** Heatmap clusters displayed the distance of all samples based on the existence of prescribed herbal species using hierarchical clustering, and network clusters illustrated these differences in Cytoscape based on ITS2 (A and B) and *trnL* (C and D) sequencing results. For heatmap (A & C), the gradient color bars mean the distance between any two samples, while the red and the blue color depicts the two extreme distances between samples. For network (B & D), each edge represents the distance of any two samples with distance less than 3.2 for ITS2 and 3.2 for *trnL*.

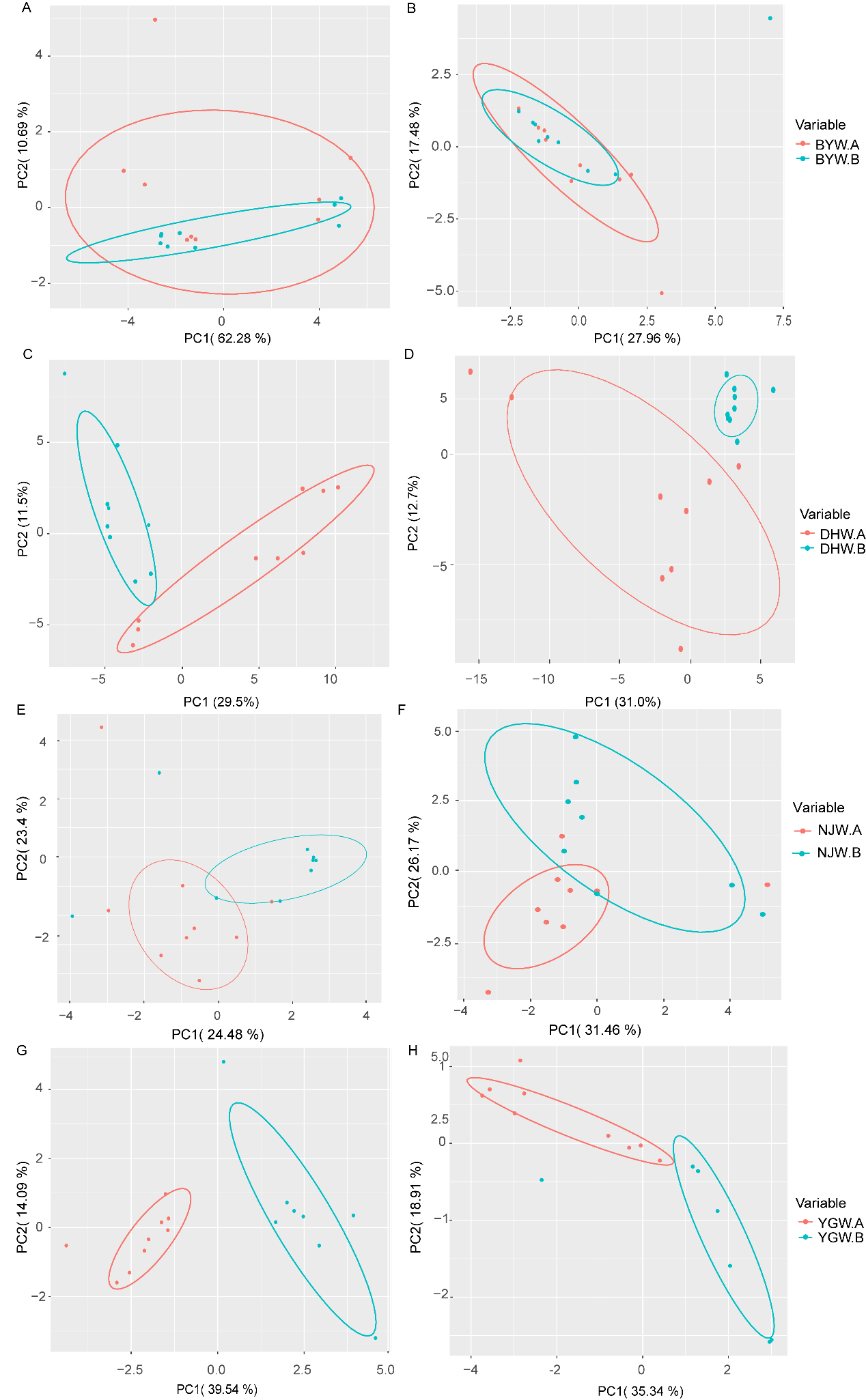

**Supplementary Figure 9. PCA analysis for assessing the consistency of the samples from intra-/inter manufacturer comparison.** (A) The samples of BYW based on ITS2; (B) The samples of BYW based on *trnL*; (C) The samples of DHW based on ITS2; (D) The samples of DHW based on *trnL*; (E) The samples of NJW based on ITS2 biomarker; (F) The samples of NJW based on *trnL* biomarker; (G) The samples of YGW based on ITS2; (H) The samples of YGW based on *trnL*.

**Supplementary Tables**

**Supplementary Table 1. The information for TCM preparations used in this study.**

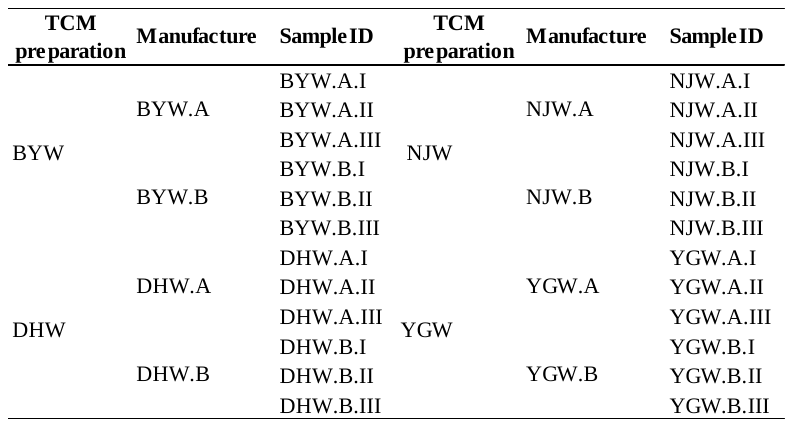

Note: “BYW.A.I” means Bazhen Yimu Wan produced by manufacturer A with batch I.

**Supplementary Table 2. ITS2 and *trnL* gene primer sequences.**

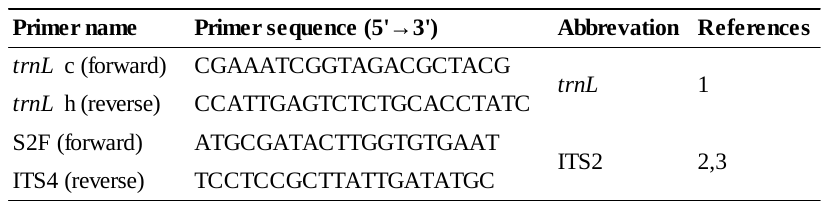

**Supplementary Table 3. The barcodes of samples used for Illimina MiSeq PE300 sequencing.**

**(a) Based on ITS2 biomarker;**

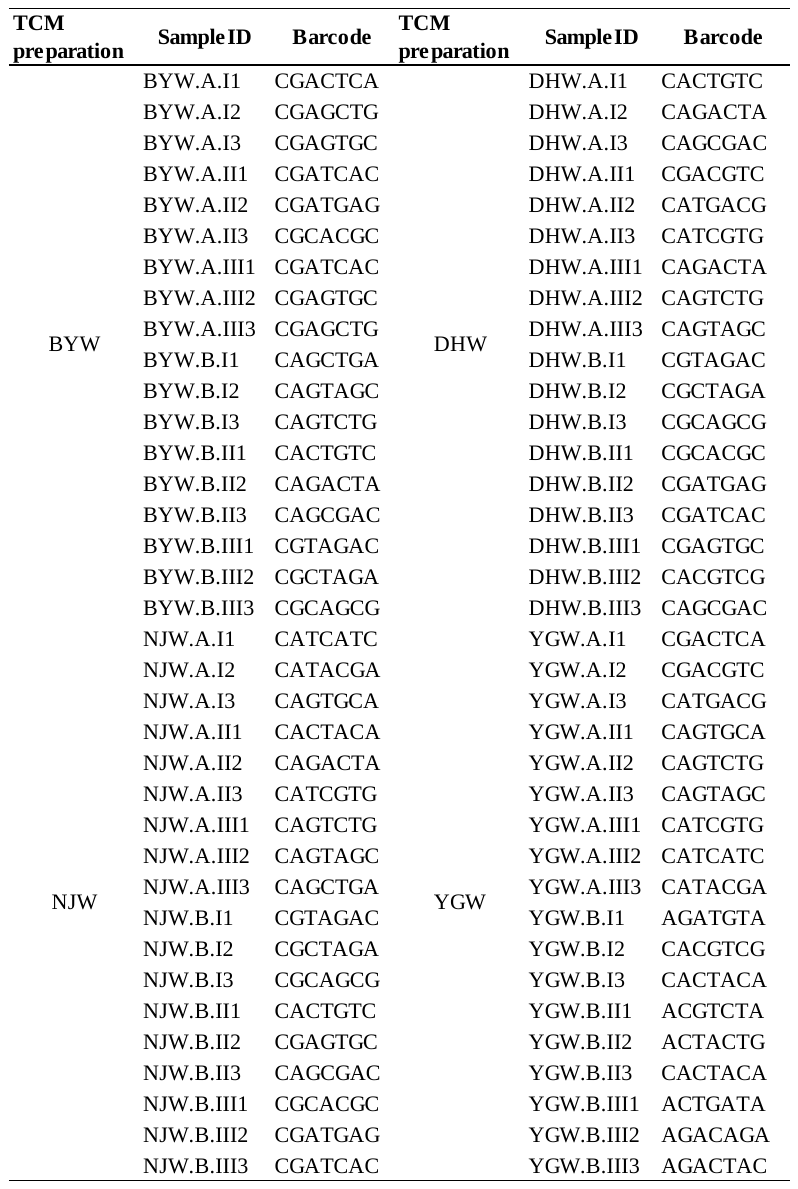

**(b) Based on *trnL* biomarker;**

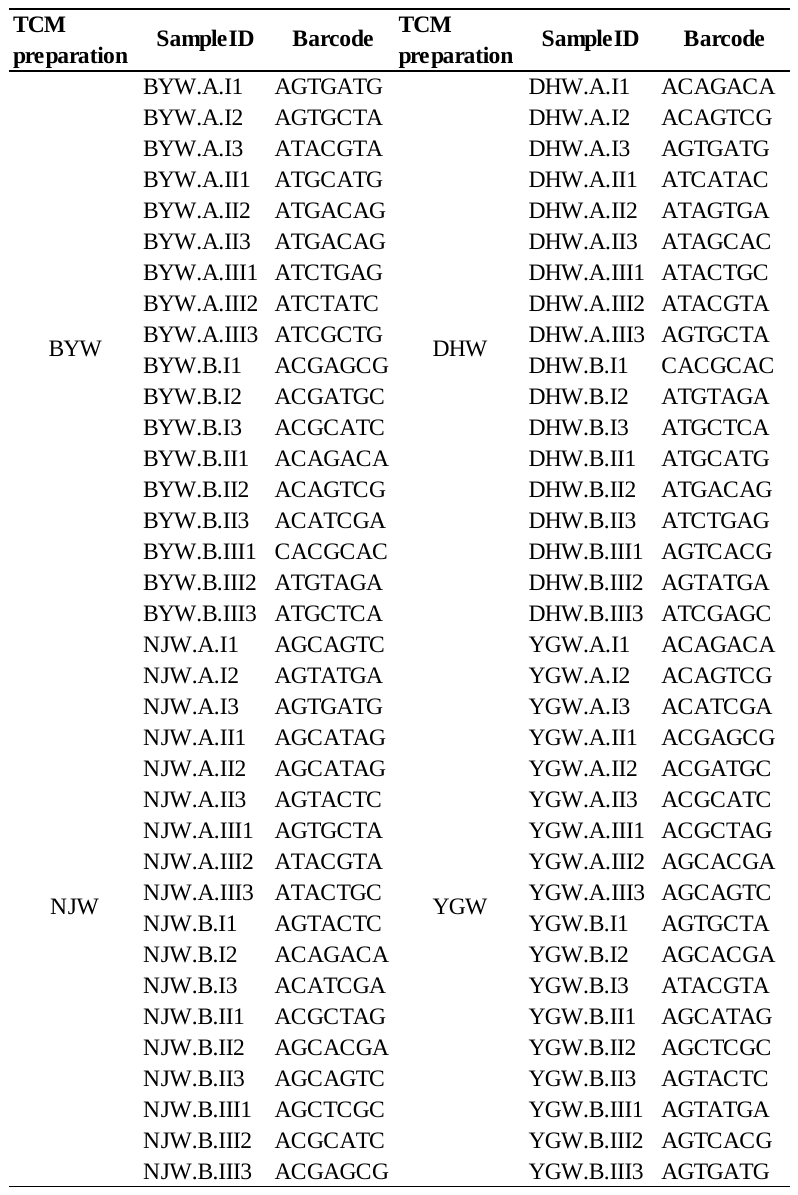

**Supplementary Table 4. The number of samples of each TCM preparation based on ITS2 and *trnL* biomarker.**

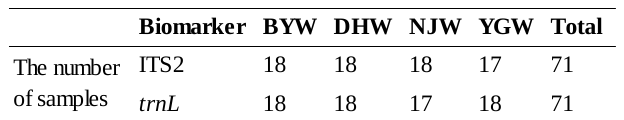

Note that one *trnL*-marked BYW sample was failed to amplify because of severe PCR inhibition, and one ITS2-marked YGW sample was failed to build library.

**Supplementary Table 5. The abbreviation of prescribed herbal materials, un-prescribed herbal materials and its corresponding prescribed herbal species of (a) Bazhen Yimu Wan (BYW), (b) Da Huoluo Wan (DHW), (c) Niuhuang Jiangya Wan (NJW).**

**(a) Bazhen Yimu Wan (BYW);**

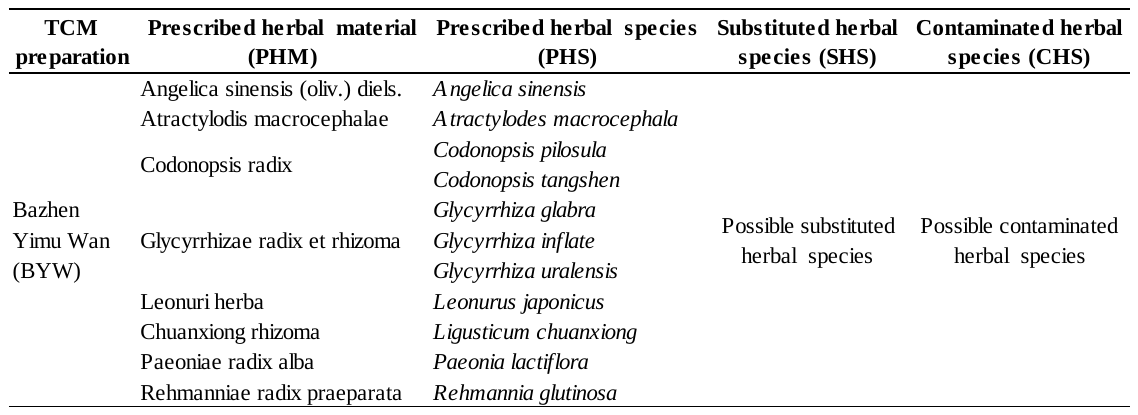

**(b) Da Huoluo Wan (DHW);**

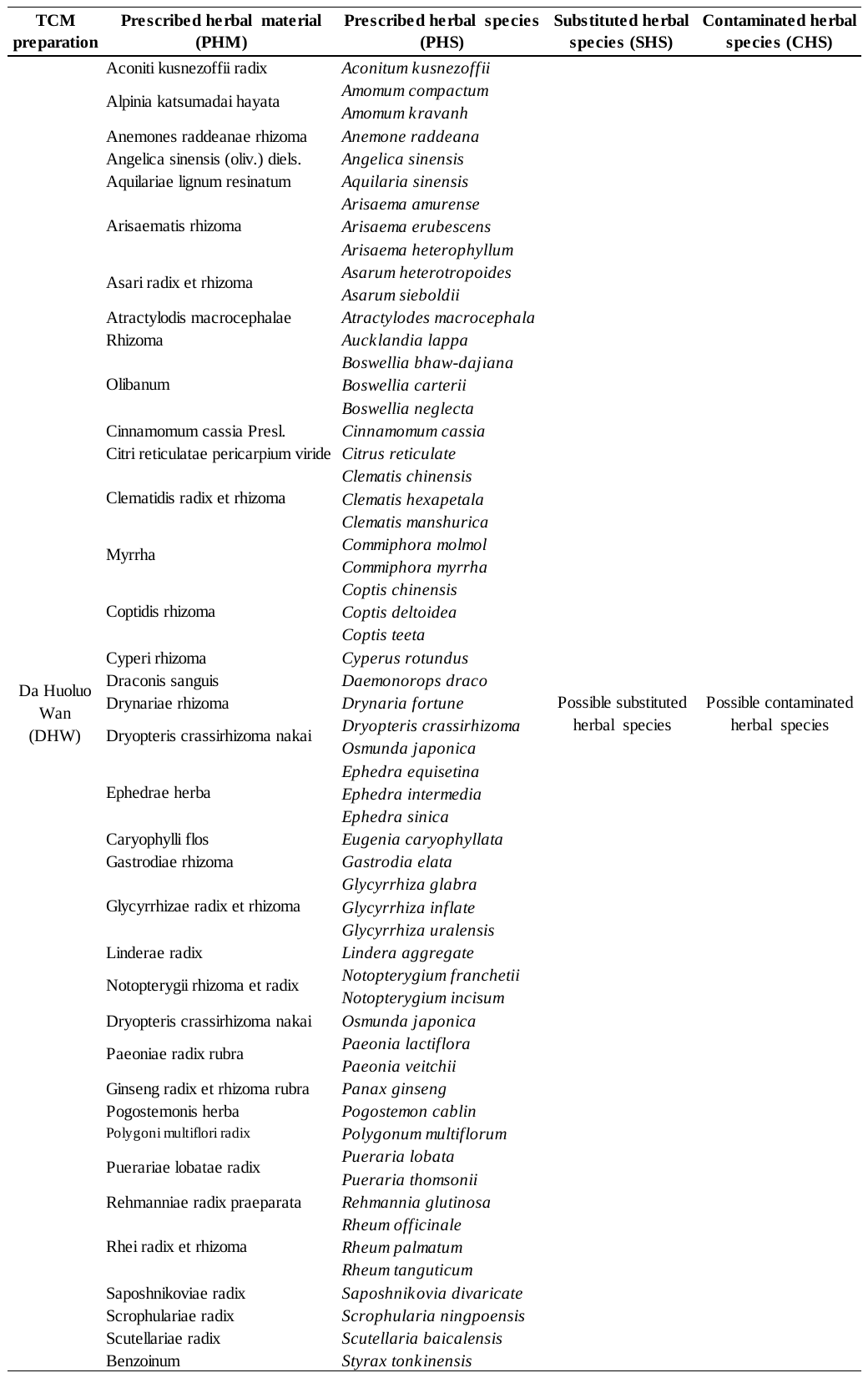

**(c) Niuhuang Jiangya Wan (NJW).**

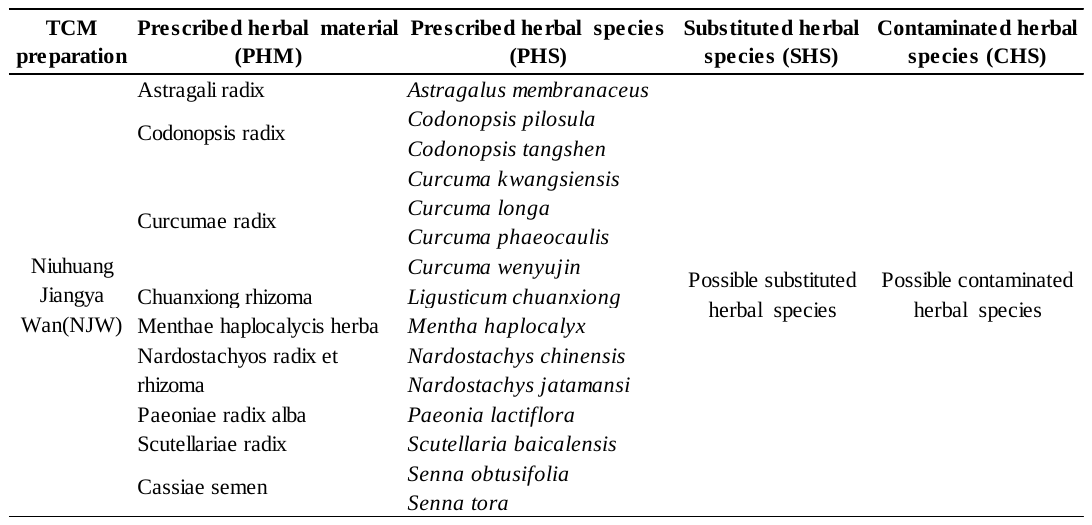

**Supplementary Table 6. The information about the animal and mineral materials for the four TCM preparations.**

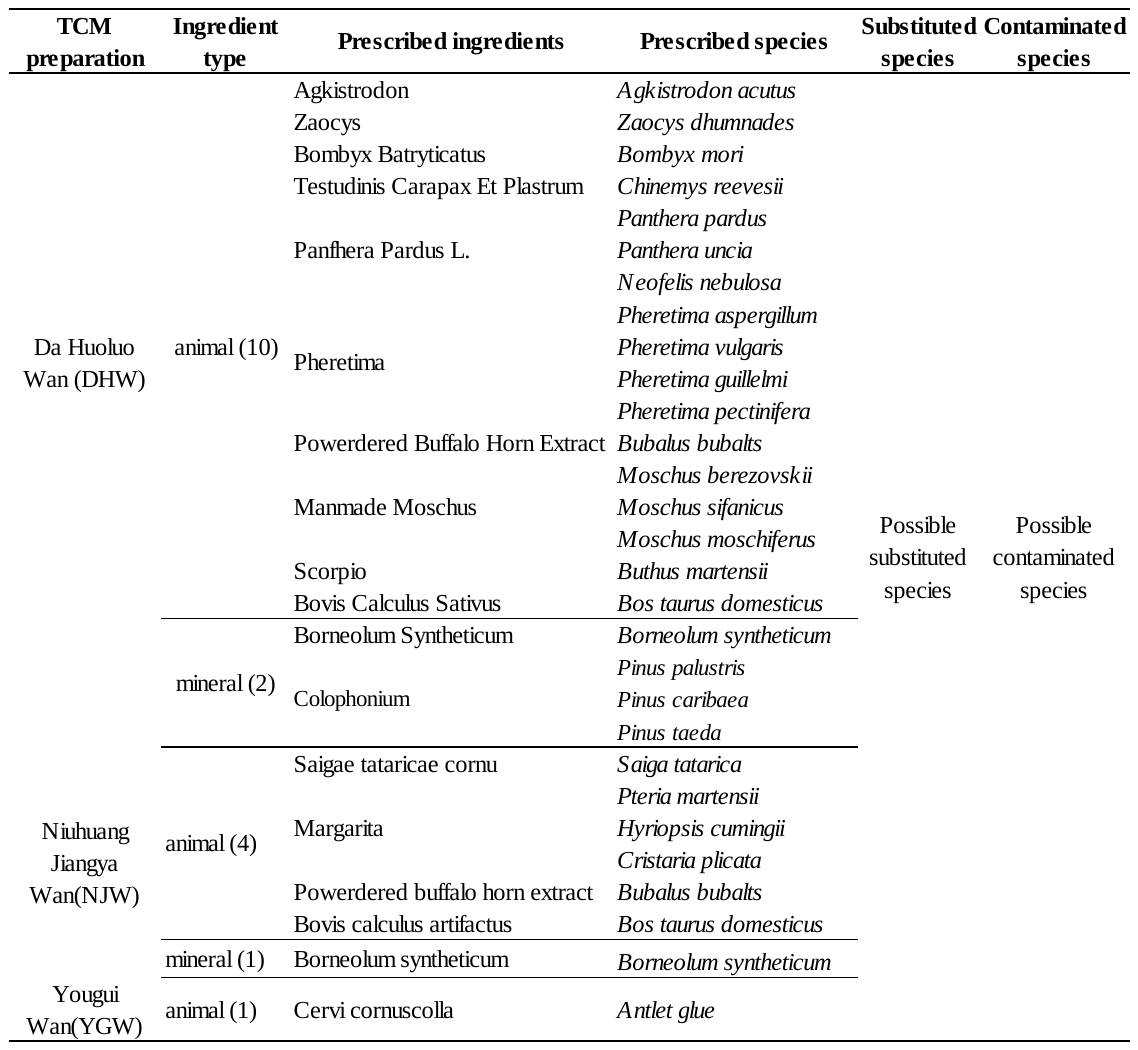

Note that Bazhen Yimu Wan (BYW) only contains herbal materials, and YGW does not contain mineral material.

**Supplementary Table 7.** **Prescribed herbal species for BYW preparation and their presence in each sample by multi-barcoding approach based on ITS2 biomarker.**

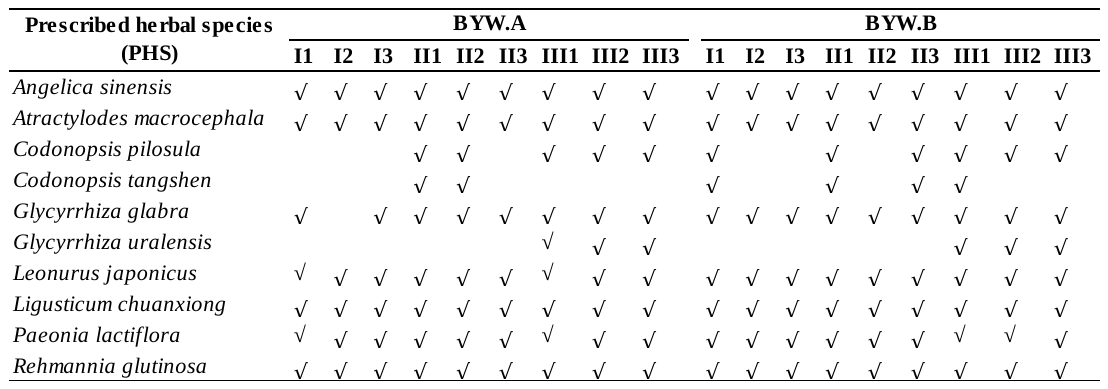

**Supplementary Table 8. Prescribed herbal species for BYW preparation and their presence in each sample by multi-barcoding approach based on *trnL* biomarker.**

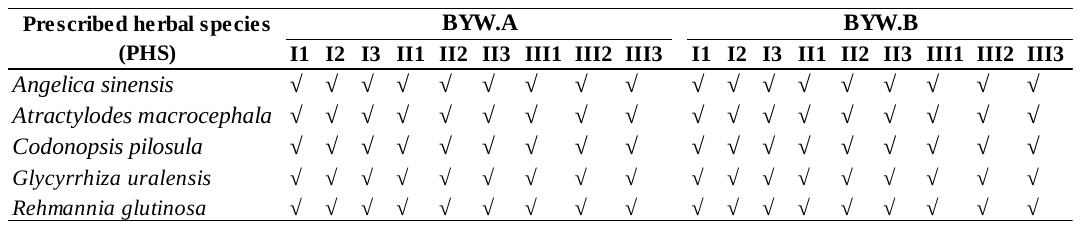

**Supplementary Table 9. Prescribed herbal species for YGW preparation and their presence in each sample by multi-barcoding approach based on ITS2 biomarker.**

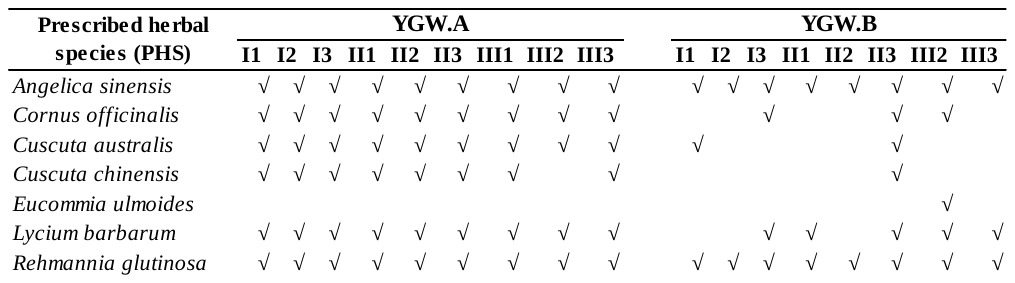

**Supplementary Table 10. Prescribed herbal species for YGW preparation and their presence in each sample by multi-barcoding approach based on *trnL* biomarker.**

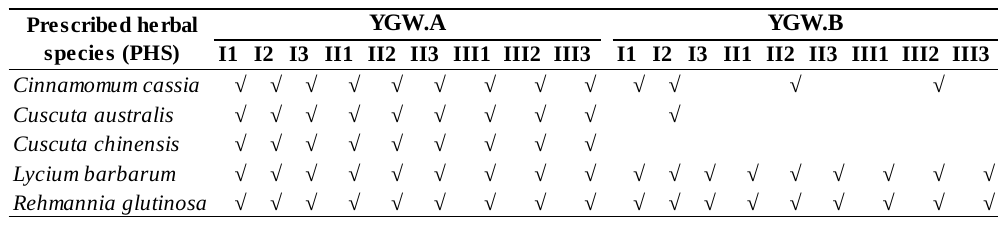
